# The proteomic landscape and temporal dynamics of mammalian gastruloid development

**DOI:** 10.1101/2024.09.05.609098

**Authors:** Riddhiman K. Garge, Valerie Lynch, Rose Fields, Silvia Casadei, Sabrina Best, Jeremy Stone, Matthew Snyder, Connor Kubo, Arata Wakimoto, Zukai Liu, Chris D. McGann, Jay Shendure, Lea M. Starita, Nobuhiko Hamazaki, Devin K. Schweppe

## Abstract

Gastrulation is the process by which the early embryo establishes a body plan and primes itself for organogenesis. As gastrulation is challenging to study *in vivo*, stem cell-derived “gastruloids” have emerged as powerful surrogates. Although transcriptomics and imaging have been applied extensively to such embryo models, the dynamics of their proteomes remains largely unknown. Here, we apply quantitative proteomics to human and mouse gastruloids at four key stages. We leverage these data to map the expression dynamics of protein complexes, and to nominate cooperative proteins. With matched transcriptome and phosphosite data, we investigate global and stage/pathway-specific discordance between the transcriptome and proteome and nominate kinase-substrate relationships based on phosphosite dynamics. Finally, we apply co-regulation network analysis to identify genes linked to the Commander complex whose perturbation leads to morphological defects in gastruloids. Altogether, our work showcases the potential of applying proteomics to embryo models to advance our understanding of mammalian development in ways challenging through transcriptomics alone.

## INTRODUCTION

Gastrulation is a crucial process in metazoan development through which the implanted blastocyst transforms into a three germ layer structure, the gastrula^1^. Ethical and practical challenges prevent us from routinely obtaining or culturing gastrula-stage human embryos, such that our understanding of human gastrulation remains limited^2,3^. Although conserved aspects of mammalian gastrulation can be studied *in vivo* in the mouse, this also suffers from practical challenges (*e.g.* opacity, limited material, and the cost of genetic manipulation). Furthermore, mouse and human gastrulation are dissimilar in many respects. Most obviously, mouse gastrula are shaped like “egg cylinders”, while human gastrula, like most other mammals, are flat discs^4^. These morphological contrasts are accompanied by differences in the expression or source of key regulators (*e.g.* FGF8, BMP4) as well as in the origins and timing of appearance of various cell types (*e.g.* primordial germ cells, extraembryonic ectoderm)^4^.

*In vitro* stem cell-derived embryo models are powerful surrogates for *in vivo* embryos, and in recent years have proliferated not only in usage but also with respect to the specific aspects of embryogenesis that are modelled^5^. One particular class of embryo models, gastruloids, are generated by first aggregating hundreds of embryonic stem cells (ESCs) and then inducing WNT signaling, which triggers axial elongation and the emergence of all three germ layers over the ensuing days^6–8^. In the presence of extracellular matrix-like scaffolding material (“Matrigel”), mouse gastruloids form morphological structures resembling their *in vivo* counterparts, including an elongated neural tube and flanking somites^8,9^. Recently, we demonstrated that early retinoic acid, together with Matrigel, yields human gastruloids with these same morphological structures, as well as advanced cell types including neural crest, neural progenitors, renal progenitors, and myocytes (“RA-gastruloids”)^10^. Importantly, gastruloids can be chemically and/or genetically manipulated, visually and/or molecularly characterized, and, owing to their ease of culturing, even grown in large numbers^7^.

Various groups, including us, have subjected time-courses of gastruloids to scRNA-seq to characterize the dynamics of the transcriptome as ESCs diversify into germ layers and cell types^11,12^. However, RNA is only the messenger. It is proteins that are the workhorses of the cell, and in the context of differentiating gastruloids, proteins that form the structures that make emerging germ layers and cell types morphologically and functionally unique. It is challenging to accurately estimate protein abundances from transcriptomics alone^13–18^. Studies report varying levels of discordance^19–23^, *e.g.* one recent study found that in human cells, transcript abundance accounted for only ∼40% of the variance in protein levels^24^. Moreover, post-translational modifications (PTMs) including phosphorylation, ubiquitination, and glycosylation vastly increase the functional diversity of cells’ proteomes to over ∼10 million proteoforms^25^, aspects of identity and function that are entirely absent from a transcriptomic census. Such PTMs are known to dynamically regulate signaling pathways that critically underpin developmental patterning and cell type specification, *e.g.* WNT, BMP and FGF^26^. However, only a handful of studies to date have attempted to characterize the proteome in early mammalian developmental contexts, and, to our knowledge, none in human post-implantation embryos or gastruloids^13,27^.

Here we applied high-throughput quantitative mass spectrometry to quantify proteins and phosphosites across four key stages of gastruloid differentiation. With these data, we map the dynamics of hundreds of known protein complexes, while also identifying additional proteins whose temporal profiles correlate with specific complexes, suggesting cooperative relationships during early development. With experimentally matched RNA-seq data, we identify pathway-specific patterns of concordance and discordance between the transcriptome and proteome during gastrulation. Extending our study to phosphorylated proteins, we mapped the dynamics of thousands of phosphosites to predict stage-specific kinase activities across gastruloid development. We observed that MAPKAPK2 plays crucial roles in gastruloid development by regulating exit from pluripotency. Finally, we leveraged temporal co-regulatory protein networks for genes associated with the Commander complex to establish essential roles for DPYSL4 and PRKACB in gastruloid development. Altogether, by focusing on the proteome and phosphoproteome in models of early gastrulation, these data and analyses lay the groundwork for closing the gap between transcriptomic vs. cellular views of early mammalian development. The data are made freely available together with a custom browser at: https://gastruloid.brotmanbaty.org/.

## RESULTS

### Quantifying the dynamic proteome from ESCs to gastruloids

We profiled the dynamics of RNA, protein, and phosphosite levels in human RA-gastruloids^10^ and conventional mouse gastruloids^9^. Specifically, we performed matched bulk RNA-seq, quantitative proteomics and quantitative phosphoproteomics on whole cell extracts from both human and mouse samples corresponding to four stages of gastruloid differentiation, including two ESC stages (“naïve” and “primed”) and two gastruloid stages (“early” and “late”) (**Fig. 1a**; **Extended Data Fig. 1a**). We focused on these four stages because they model pre-implantation, post-implantation, post-symmetry breaking, and anterior-posterior (A-P) elongation/patterning, respectively. For human primed ESCs, we analyzed two cell lines (H9, RUES2-GLR) to assess inter-cell-line variation^28,29^ (**Fig. 1b**). As such, we analyzed nine sample types altogether—four mouse (naïve ESCs, primed ESCs, early gastruloids, late gastruloids) and five human (naïve H9 ESCs, primed H9 ESCs, primed RUES2-GLR ESCs, early RUES2-GLR gastruloids, late RUES2-GLR gastruloids)—in biological duplicate (transcriptomics) or triplicate (proteomics, phosphoproteomics) (**Extended Data Fig. 1b-c**).

**Figure 1.**
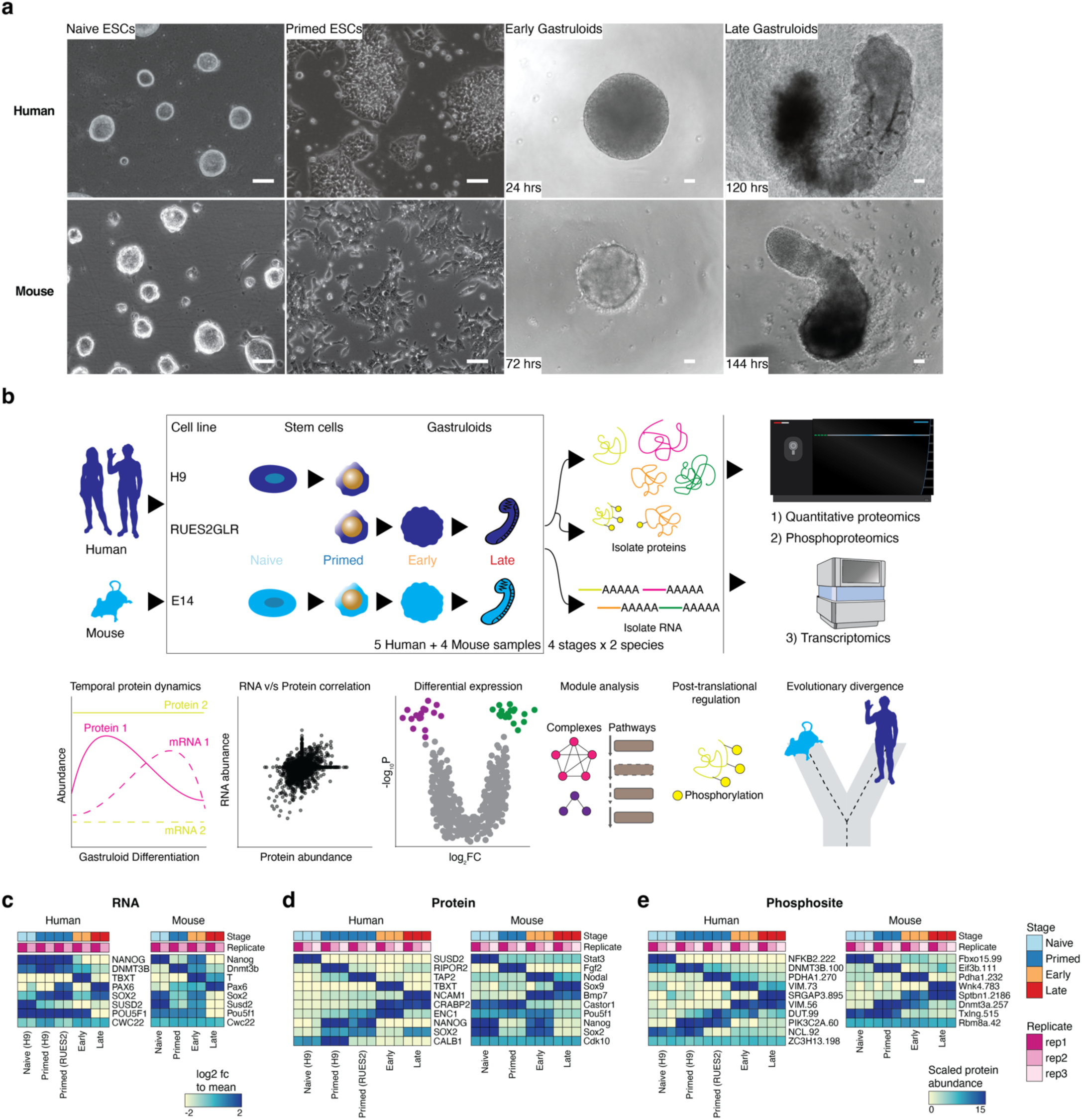
Quantifying the dynamic proteome from ESCs to gastruloids. **(a)** Representative brightfield images of human RA-gastruloids and mouse gastruloids imaged over the course of their development. Scale bar: 10 μm. **(b)** Multi-omics profiling workflow. We sampled two human cell lines (H9 and RUES2-GLR) and one mouse cell line (E14) at the indicated stages. **(c-e)** Representative heatmaps depicting the temporal dynamics of RNAs **(c)**, proteins **(d)**, or phosphosites **(e)** for selected developmental marker transcripts, proteins or PTMs, respectively, across replicates and stages for both human and mouse. Color scale for RNAs indicates log2-fold change relative to the row mean. Color scale for protein and phosphorylation data indicates scaled TMT abundance.

To assess data quality, we calculated pairwise correlations between biological replicates of each sample type and confirmed high reproducibility for each data type (RNA: *r* > 0.98; protein: *r* > 0.93; phosphosite: *r* > 0.97; **Extended Data Fig. 1c-e**). Consistent with that, replicates for each data type were tightly grouped by Principal Components Analysis (PCA) (**Extended Data Fig. 1f**). In human, for all three data types, PC1 separated naïve H9 ESCs from other samples (RNA: 43%; protein: 45%; phosphosite: 52% of variance explained) while PC2 broadly correlated with developmental progression (RNA: 26%; protein: 34%; phosphosite: 21% of variance explained). In mouse, for all three data types, PC1 separated late gastruloids from other samples (RNA: 56%; protein: 50%; phosphosite: 44% of variance explained), while PC2 once again resolved developmental progression (RNA: 26%; protein: 31%; phosphosite: 33% of variance explained) (**Extended Data Fig. 1f**).

Across all replicates of all stages, we detected and quantified 7,352 human and 8,699 mouse proteins (**Extended Data Fig. 1b**; **Supplementary Table 1**). To gauge the depth of proteome sampling, we mapped these proteins onto the Human Protein Atlas^30^, and found all 34 annotated subcellular locations to be represented (**Extended Data Fig. 1g**). To confirm that we are capturing developmental transitions, we searched for stage-specific markers at both the RNA and protein levels in each species. Results were generally consistent with expectation, *e.g.* in human samples, classic pluripotent markers *NANOG* and *POU5F1* were highly expressed in ESCs, while *TBXT*, a marker of gastrulation and/or mesendoderm differentiation^31^, and *PAX6*, associated with neural tube differentiation^32^, were upregulated in early and late stage gastruloids, respectively (**Fig. 1c**). Many detected proteins also exhibited stage specificity. For example, again focusing on human samples, SUSD2, a cell-surface marker for the naïve epiblast^33^, was only detected in naïve H9 cells, while TBXT and NCAM1 were specific to early and late stage gastruloids, respectively (**Fig. 1d**). Interestingly, CRAPBP2, a retinoic acid binding protein^34^, was detected only in human samples after addition of retinoic acid into the culture media^35^. For some markers, we observe consistent dynamics for mRNA and protein abundance, *e.g.* mouse *Sox2*/Sox2 (**Fig. 1c,d**). We observed that the pluripotency marker SOX2’s protein levels in early stage gastruloids display a marked reduction before rising back up in late gastruloids. We validated our observations and multi-omic approach by temporally profiling SOX2 endogenously tagged with mCitrine and found that this pattern was driven by neural cell populations (neural progenitors, neural crest, and neural tube cell types) arising 48 hours after induction (**Fig. 1c,d**; **Extended Data Fig. 1h**). Using sample multiplexed quantitative phosphoproteomics^17,36^, to map temporal dynamics of human and mouse phosphosignaling (**Fig. 1e**) and mapped their temporal dynamics across the four stages. We observed decreased phosphorylation of the methyltransferases DNMT3B (Ser100, human) and Dnmt3a (Thr257, mouse) during gastruloid development relating gastruloid cellular signaling to previous reports of DNA hypomethylation in ground state pluripotency and increased methylase activity during differentiation^26,37–41^. Compared to recent mouse gastruloid proteomics datasets, we quantified 3,290 more mouse proteins (65% increase) and 2,303 more homologous human proteins (46% increase). High overlap with murine embryonic proteome datasets confirmed that we sampled biologically relevant temporal protein changes (**Extended Data Fig. 2c,d**). The increased depth of the proteome sampled over the course of gastruloid differentiation enabled temporal coregulatory analysis at the level of proteins, complexes, and phosphosignaling (**Extended Data Fig. 2a,b**). Taken together, these data along with the dedicated web application enable exploration and new understanding of the temporal dynamics of RNA, protein, and cell signaling models of human and mouse gastrulation.

**Extended Data Figure 1.**
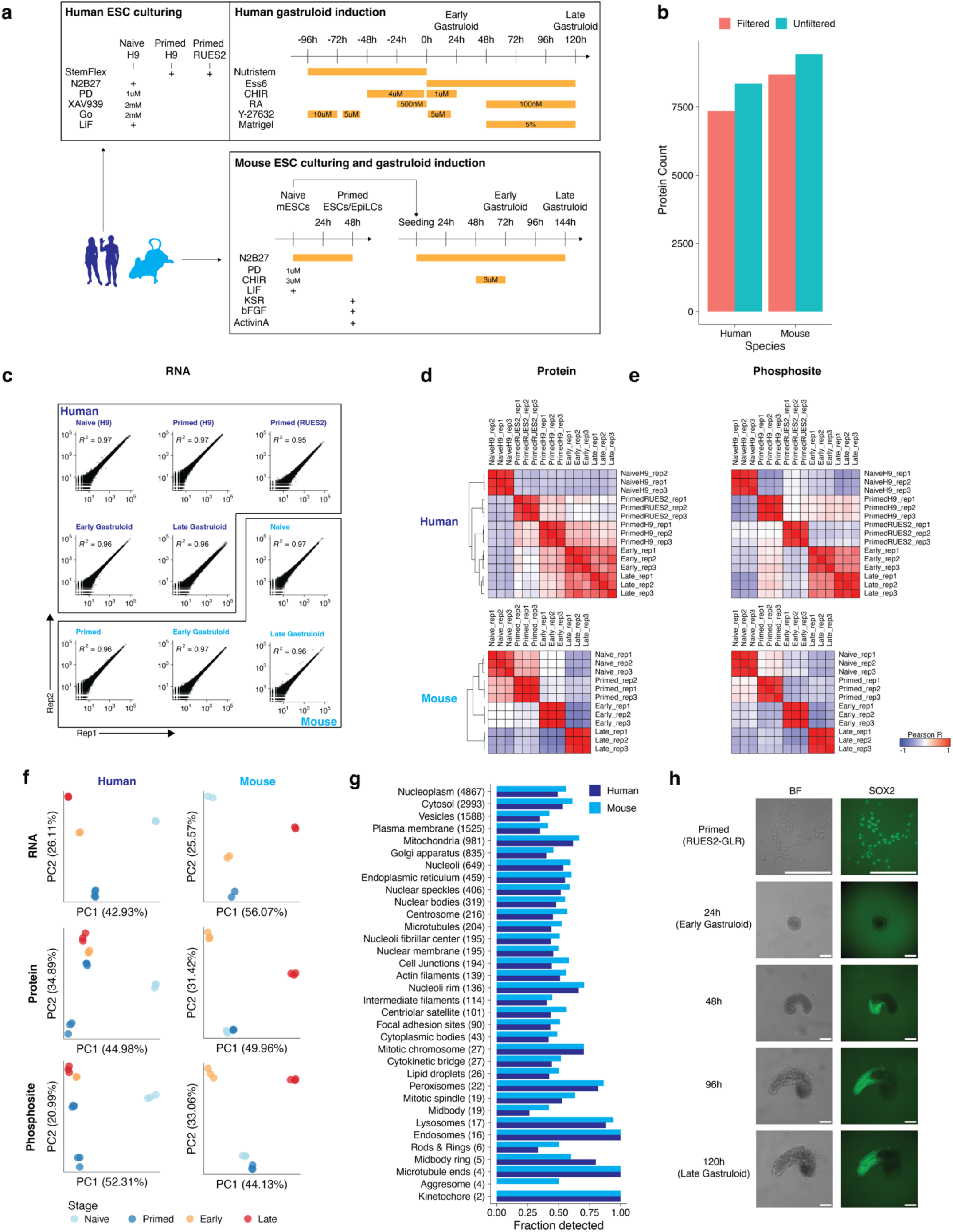
Mapping the dynamics of gastruloid development using multi-omics. **(a)** Timeline and conditions for human and mouse ESC culturing and gastruloid induction. **(b)** Total numbers of proteins quantified across all human or mouse samples. Protein identifications were filtered to a 1% FDR and required summed TMTpro reporter ion signal-to-noise ratios >100 for quantitation. **(c)** Scatterplots comparing the RNA counts between biological replicates for each sample. **(d,e)** All-by-all sample similarity matrices of pairwise Pearson correlation coefficients (r_Pearson_) calculated from summed protein **(d)** or phosphosite intensities **(e)** across human (top) or mouse (bottom) samples. **(f)** PCA plots of PC1 vs. PC2 using RNA (top), protein (middle) or phosphosite (bottom) data across human (left) or mouse (right) samples. **(g)** Fraction of proteins assigned to each of 34 subcellular localizations by the Human Protein Atlas that were successfully detected here. Numbers within brackets indicate the total numbers of proteins within each class shown. **(h)** Representative images highlighting the morphology (left) and the SOX2-mCitrine expression across the stages of gastruloid development. Scale bar: 200 μm.

**Extended Data Figure 2.**
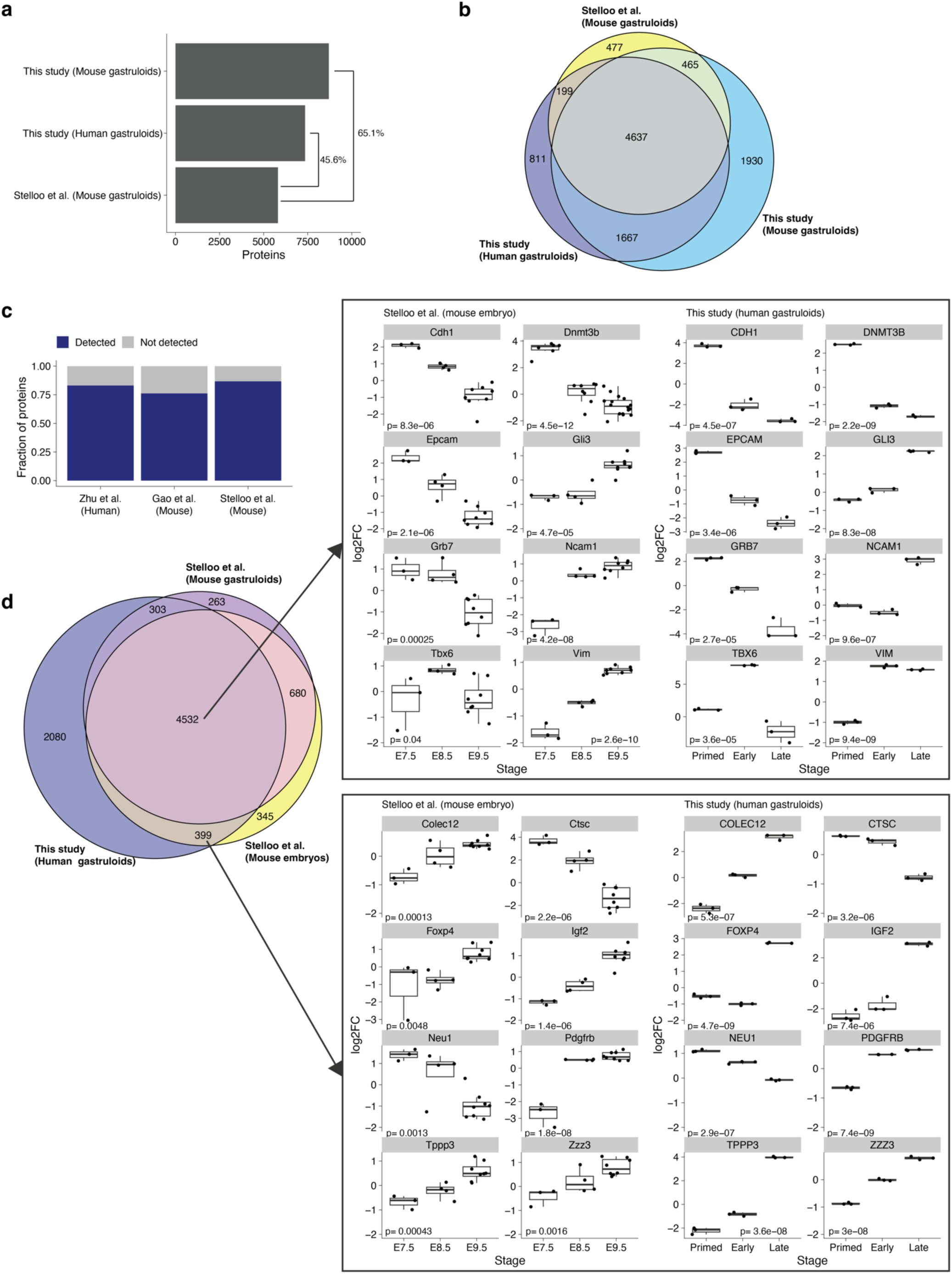
**(a)** Comparison of this study with other Stelloo et al., **(b)** Venn diagram representing the fraction of additional proteins detected in our datasets with Stelloo et al., **(c)** Fraction of proteins detected in this study compared to existing human and mouse embryo proteomics datasets. **(d)** Venn diagram of overlap of proteins with aforementioned datasets (right) and temporal trends in embryo datasets recapitulated by this study.

### Time-resolved proteomics reveals biologically coherent shifts across gastruloid development

To identify sets of proteins with similar temporal dynamics, we merged the human and mouse proteomic datasets by orthology and subjected them to hierarchical clustering (**Fig. 2a**). Focusing on 10 clusters, *i.e.* sets of proteins that exhibit similar dynamics across both species, we assessed Gene Ontology (GO) enrichments^42^. Seven of the 10 clusters returned significantly enriched biological processes, *i.e.* cell division and DNA repair (cluster 1), mitochondria and aerobic respiration (cluster 2), RNA biogenesis (cluster 3), cilia and pattern specification (cluster 4), small molecule metabolism (cluster 6), extracellular matrix organization (cluster 7) and tube development (cluster 8) (**Fig. 2b**; **Supplementary Table 2**). These enrichments suggest that the abundance of proteins that underlie these biological processes are coordinated during gastrulation.

**Figure 2.**
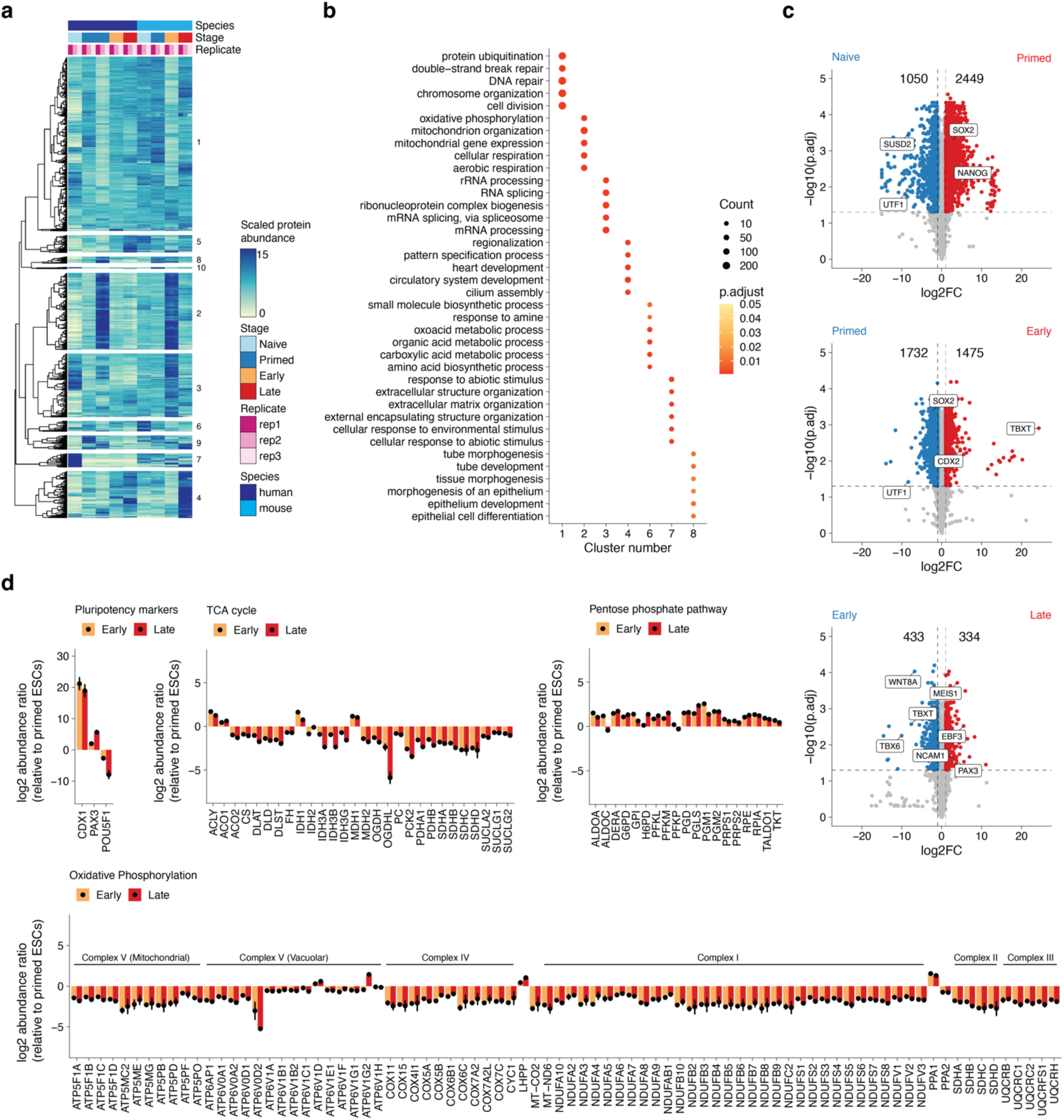
Time-resolved proteomics reveals biologically coherent shifts across gastruloid development. **(a)** Heatmap depicting the temporal dynamics of protein expression across human and mouse gastruloid differentiation samples and replicates. **(b)** Dotplot indicating the top Gene Ontology (GO) terms enrichment across clusters. Clusters 5, 9, and 10 did not have significantly enriched GO terms. Color scale indicates the Benjamini Hochberg (BH) adjusted p-values. Size of dots corresponds to the number of proteins associated with a particular GO term. **(c)** Volcano plots of the protein expression changes across consecutive stages of human gastruloid differentiation, where x-axis represents the log2 fold change between two adjacent timepoints and y-axis represents the negative log_10_ of the Benjamini-Hochberg-adjusted p-value. **(d)** The log_2_ protein abundance ratio of early (yellow) or late (red) gastruloids compared to primed human ESCs (RUES2-GLR) for proteins associated with pluripotency and central metabolism including TCA cycle, pentose phosphate pathway and oxidative phosphorylation. Mean abundance ratios are indicated with dots and error bars represent the standard deviation.

To identify the changes in protein abundance that may underlie specific transitions in gastruloid development, we performed differential expression analysis across adjacent timepoints in each species, which identified thousands of differentially expressed proteins (DEPs) (**Extended Data Fig. 3b**). For example, when comparing naïve and primed states of pluripotency within human H9 cells, we identified 3,499 DEPs. Among these was SUSD2, whose expression marks pre-implantation epiblasts in human blastocysts, which was detected only in the naïve state, as well as SOX2 and NANOG, which were enriched in the primed state (**Fig. 3c**). GO analysis of naïve vs. primed DEPs found that naïve cells were enriched for proteins involved in extracellular matrix (ECM) organization, while primed cells were enriched for proteins involved in nucleotide metabolism (**Extended Data Fig. 3d**). In comparing primed RUES2-GLR ESCs to early human RA-gastruloids, we identified 3,207 DEPs, including SOX2 enrichment in primed ESCs, and TBXT and CDX2 enrichment in early human RA-gastruloids. Upon GO analysis, DEPs upregulated in early gastruloids mapped to actin filament organization and cytoskeletal processes, while DEPs downregulated mapped to mitochondrial processes (**Extended Data Fig. 3d**). In comparing early vs. late human RA-gastruloids, we identified 767 DEPs, including downregulation of TBXT, caudal axial progenitors marker WNT8A, and presomitic mesoderm marker TBX6, and upregulation of markers of advanced cell types including PAX3 (dorsal somites and neural tube), SOX1 & SOX2 (neural tube) and cardiomyocytes (MEIS1) (**Fig. 2c**).

**Figure 3.**
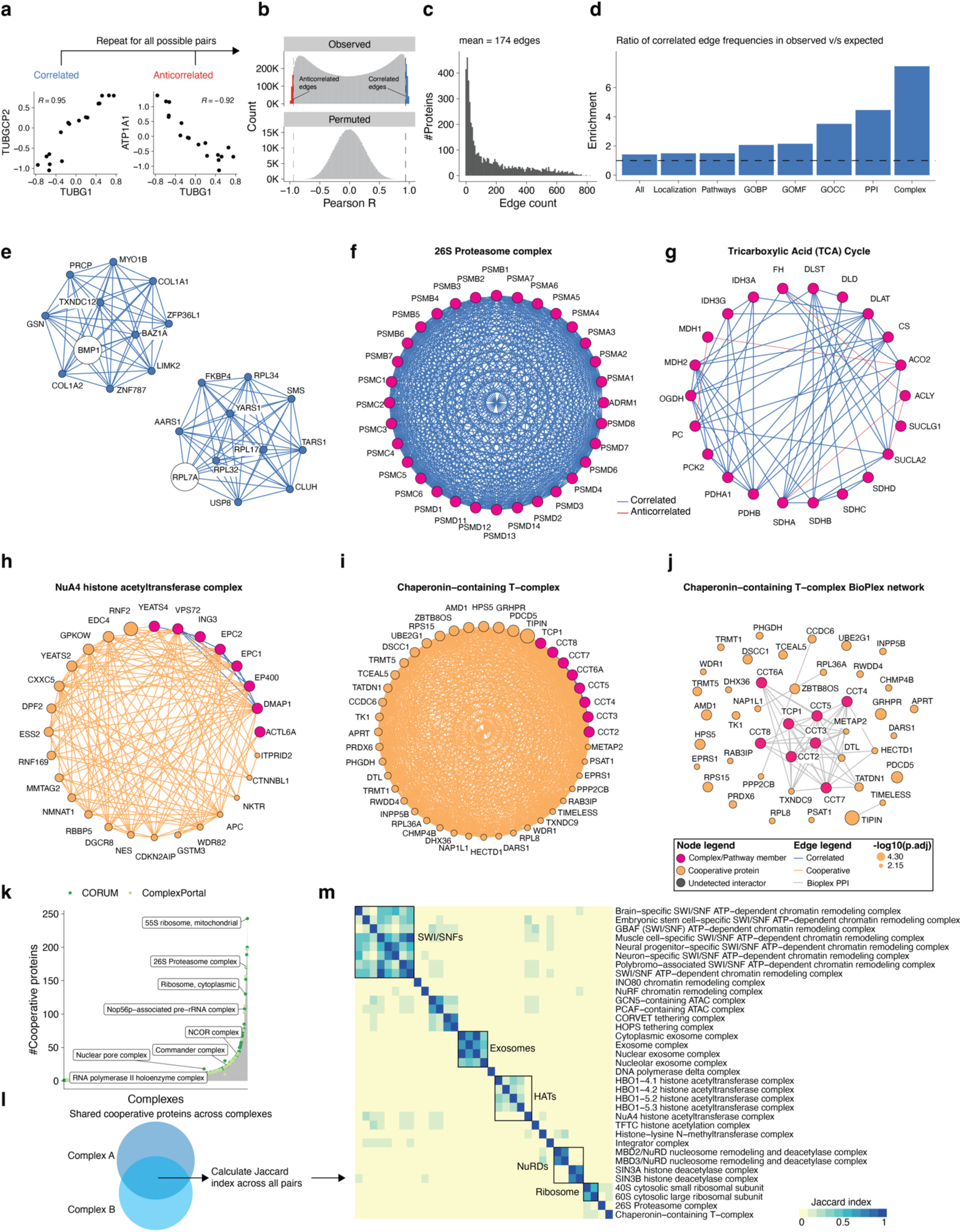
Co-regulation analysis maps cooperative protein associations to known protein complexes and pathways. (a) Scatterplots comparing abundances across selected protein pairs across samples. **(b)** Distribution of r_Pearson_ based on observed (top) and permuted (bottom) data. Observed distribution was obtained by calculating r_Pearson_ across all possible protein-protein pairs. Permuted distributions were generated by randomly sampling 50,000 protein pairs after randomly shuffling their respective timepoints 10 times each prior to calculating r_Pearson_. Colors indicate strongly correlated (>=0.95; blue) or anticorrelated (<=-0.95; red) edges. **(c)** Distribution of protein edge counts across the trimmed correlation network. On average, each protein in the network participated in 174 edges. **(d)** Ratio of enrichment for the annotated edges in the correlation network (“observed network”) compared to the expected edge annotation frequencies across Gene Ontology biological process (GOBP), cellular component (GOCC), molecular function (GOMF), localization, pathways, protein-protein interactions (BioPlex) or protein complexes. Specifically, we calculated the enrichment for annotated edges as the fraction of annotated edges per category in the observed correlation network divided by the fraction of annotated edges among all possible edges involving the 5,227 proteins in the correlation network. The expected frequency of annotated edges was calculated by generating all possible pairs from 5,227 human proteins (Uniprot, 07/2024) and computing the number of pairs explained by each functional category.**(e)** Network analysis identifies known associations between proteins for BMP1 and RPL7A. **(F-G**) Network structure of the **(f)** 26S proteasome and **(g)** Citric Acid cycle pathway. Magenta nodes indicate known complex members annotated either from CORUM or EMBL ComplexPortal for protein complexes, or from BioCarta, KEGG, Protein Interaction Database (PID), Reactome, and WikiPathways (WP) for biochemical pathways. Blue edges indicate positive correlations between nodes while red edges indicate anticorrelations. **(H-I)** Cooperative proteins are highly correlated with members of established protein complexes including: **(h)** NuA4 chromatin remodeling complex and **(i)** Chaperonin-containing T (TRiC/CCT) complex. Magenta nodes indicate subunits of a given complex, while orange nodes indicate cooperative proteins *i.e.* proteins with correlated profiles to proteins constituting a particular protein complex. Cooperative node sizes indicate the negative log10 of the BH-adjusted p-value after computing significance from Fisher’s exact tests to determine cooperative association of a protein to a particular module. Blue edges indicate correlated edges while orange edges link cooperative proteins to members of a particular module. **(j)** Bioplex interaction network of the TRiC/CCT complex. Orange nodes are cooperative proteins with correlated profiles to proteins found in the TRiC/CCT complex. Gray edges indicate BioPlex evidence. **(k)** Histogram of protein complexes (x-axis) and their respective numbers of cooperative proteins (y-axis). **(l)** Heuristic to identify shared cooperative proteins between complexes. **(m)** Heatmap depicting a subset of shared cooperative proteins across manually curated EMBL ComplexPortal protein complexes namely exosomes, SWI/SNFs, ATAC remodelers, nucleosome remodelers (NuRDs), and histone acetyltransferase (HAT) and deacetylase (HDAC) complexes. Heatmap colored by Jaccard similarity coefficients calculated from overlapping sets of cooperative proteins between protein complex pairs and clustered using euclidean distances with average linkage.

To identify the cell types driving our bulk proteomic observations, we compared our proteomics dataset with existing scRNA-seq data profiling gastruloid development^10^. We focused on seven proteins that showed characteristic upregulation in early gastruloids (TBXT, WNT8A, TBX6, APLNR), late gastruloids (SOX2, PAX3), or both (NEBL) (**Extended Data Fig. 5a**). Mapping these genes onto the scRNA-seq data revealed distinct patterns of cell type–specific expression. In early gastruloids, TBXT was predominantly expressed in NMP and axial mesoderm, while WNT8A and TBX6 were enriched in NMP, nascent mesoderm, and primitive streak populations (**Extended Data Fig. 5b,c**). APLNR, known for its critical role in mesoderm development^43^, showed broad expression across mesodermal lineages, suggesting that both nascent and emergent mesoderm populations contribute to its bulk protein profile. In late gastruloids, SOX2 was specifically expressed in neural populations, including neural progenitors and neural tube cells, with low expression in earlier stages. PAX3 expression was primarily driven by neural and somite populations. Interestingly, NEBL protein levels in late gastruloids tended to reflect expression in cardiac cell populations, whereas in early gastruloids it lacked clear cell type specificity (**Extended Data Fig. 5b,c**). These observations suggest that our proteomic profiling was capable of capturing both cell type-specific and broad expression patterns.

We also compared H9 vs. RUES2-GLR human primed ESCs and detected 3,047 DEPs (**Extended Data Fig. 3b**). While both cell lines expressed detectable levels of characteristic primed ESC markers (*e.g.* SOX2, NANOG), DEPs largely mapped to mitochondrial processes (respiration, oxidative phosphorylation), which are upregulated in primed RUES2-GLR relative to primed H9 ESCs. Conversely, DEPs upregulated in primed H9 ESCs were enriched for cytoskeletal processes and translation (**Extended Data Fig. 3c**). This comparison reinforces the view that there are substantial differences between these widely used human ESC lines^44^.

As such, the proteomes of primed RUES2-GLR ESCs were highly enriched for mitochondrial processes, relative to both primed H9 counterparts as well as RUES2-GLR-derived early RA-gastruloids (**Extended Data. Fig. 3c,d**), with the latter suggesting that these processes are downregulated over the course of gastruloid differentiation. We sought to ask if this downregulation was specific to a subset of mitochondrially mediated metabolic pathways, as opposed to being more general. For this, we compared primed human ESCs vs. early and late RA-gastruloids (all RUES2-GLR-derived) with respect to individual proteins, broken down by pathway. Intriguingly, we observed highly consistent levels of downregulation of mitochondrial proteins involved in the TCA cycle and oxidative phosphorylation, and upregulation of proteins involved in the pentose phosphate pathway (**Fig. 2d**). Within oxidative phosphorylation, this consistency extended to individual protein complexes, *e.g.* the levels of vacuolar subunits of the ATPase complex remained relatively stable unlike their mitochondrial counterparts (**Fig. 2d**). These observations suggest that shifts in the levels of mitochondrial machinery are highly coordinated during gastruloid differentiation, consistent with previous studies of the remodeling of metabolic complexes across multiple organ systems during mammalian aging^17^. Downregulation of mitochondrial activity was also observed in H9 early gastruloids despite lower levels of OxPhos protein levels in the H9 primed ESCs (**Extended Fig. 3e,f**).

Upon extending such analyses to the mouse data, we observed similar numbers of DEPs (**Extended Data Fig. 3e**), as well as stage-specific patterns that broadly matched expectation, *e.g.* pluripotency markers Sox2 and Nanog highly expressed in naïve mESCs compared to their primed counterparts. Similarly, when comparing primed mESCs vs. early mouse gastruloids, we observed upregulation of the mesenchymal cell marker Bmp7, and in comparing early vs. late mouse gastruloids, upregulation of endoderm marker Sox17, in the more differentiated sample. To systematically analyze conserved protein expression dynamics, we compared fold-changes across pairwise stage transitions for orthologous human and mouse proteins. While we observed modest positive correlation in the naïve to primed (r_Pearson_ = 0.17) and early to late transitions (r_Pearson_ = 0.5), there was strong anticorrelation in the primed to early transition (r_Pearson_ = -0.8). However, this anticorrelation appears to be driven by the aforereferenced elevated levels of mitochondrial proteins in primed RUES2-GLR ESCs, *i.e.* the metabolic state of primed human RUES2-GLR ESCs is better matched to that of early mouse gastruloids than that of mouse primed ESCs (**Extended Data Fig. 3f**). On comparing our mouse data with a recent proteomic study in mouse gastruloids^45^, we also observed a pronounced increase in the abundance of oxidative phosphorylation proteins at the 72 hour stage. These observations recapitulate previously observed temporal trends in mouse gastruloids and the downregulation of oxidative phosphorylation is observed irrespective of starting cell number and species specific protocols (**Extended Data Fig. 4**).

**Extended Data Figure 3.**
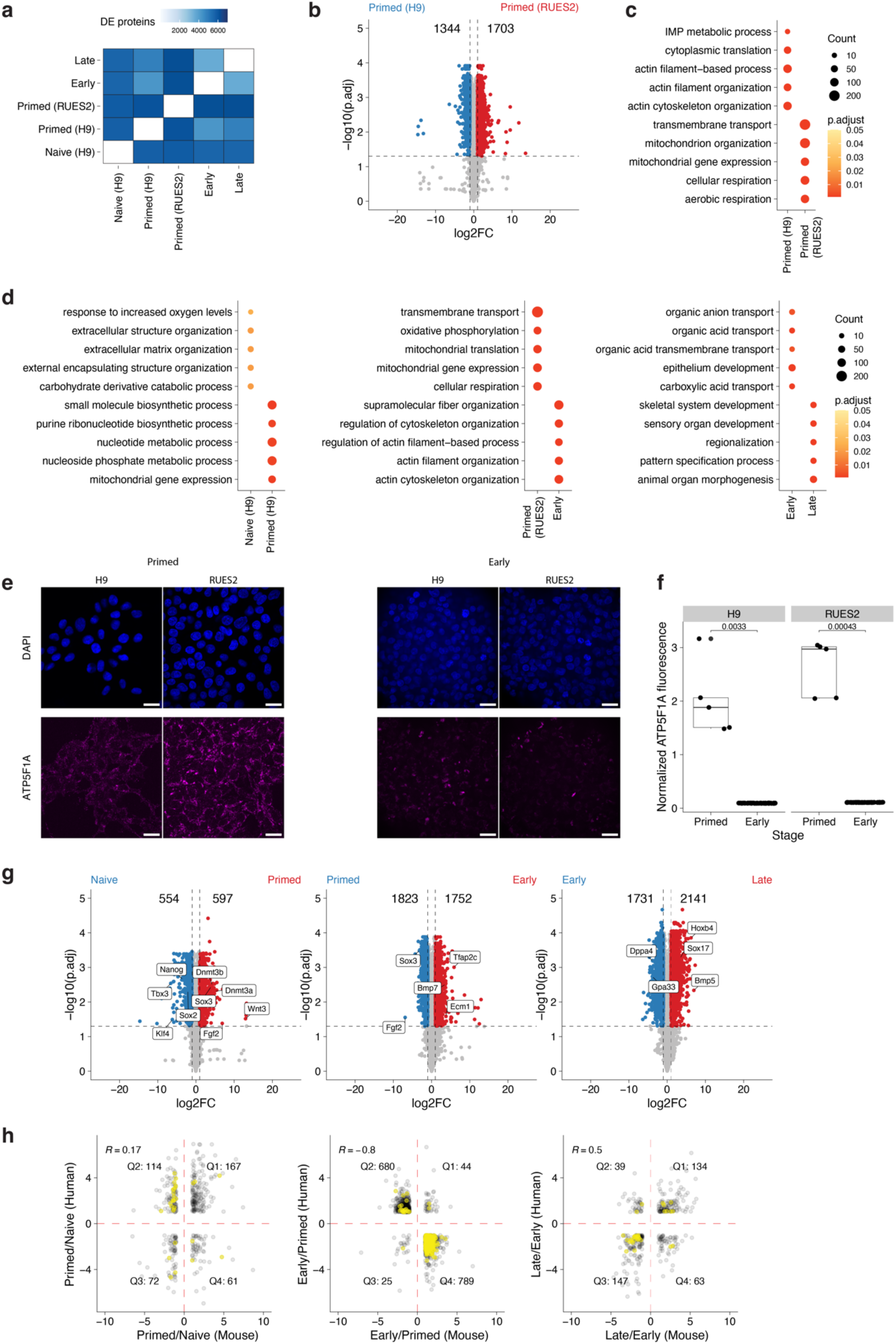
Mapping differentially expressed biological processes across gastruloid development. **(a)** Heatmap indicating the number of differentially expressed proteins (DEPs) between pairs of human samples. DEPs filtered to an absolute log_2_ fold change >= 1 and BH-adjusted p-value < 0.05. **(b)** Volcano plot depicting the DEPs between primed H9 vs. primed RUES2-GLR ESCs. **(c)** Dot plot indicating the GO terms enriched in DEPs between primed H9 vs. primed RUES2-GLR ESCs. **(d)** Dot plots indicating the GO terms enriched in DEPs between adjacent stages of human samples. Color scales for dot plots indicate the BH-adjusted p-value and sizes of dots indicate the number of genes detected within each term. **(e)** Representative images of ATP5F1A fluorescence in RUES2 and H9 primed ESCs (left) and early gastruloids (right). Blue channel indicates DAPI, Scale bar: 25 µm. **(f)** Boxplots of normalized ATP5F1A immunofluorescence intensity **(g)** Volcano plots depicting the DEPs between adjacent stages of mouse samples, where x-axis represents the log_2_ fold change between two adjacent timepoints and y-axis represents the negative log_10_ of the BH-adjusted p-value. **(h)** Scatter plots comparing mouse and human proteomes across adjacent stages. Comparisons were filtered to proteins with an absolute log_2_ fold change >= 1 across both species. Mitochondrial proteins highlighted in yellow.

**Extended Data Figure 4.**
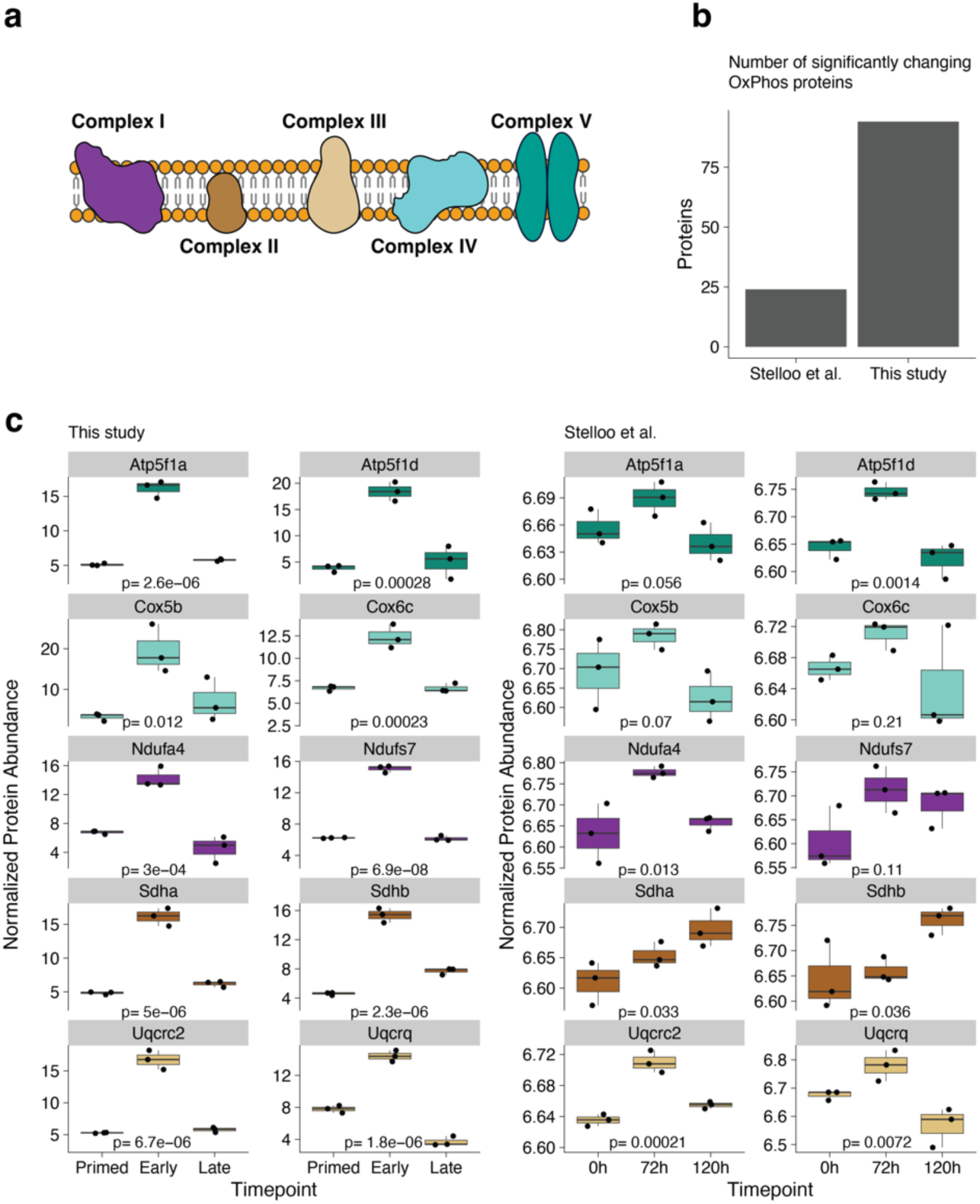
**(a)** Schematic of Oxidative Phosphorylation complexes. **(b)** Number of significantly changing OxPhos proteins in this study and Stelloo et al^45^. **(c)** Comparison of the temporal dynamics of OxPhos proteins in mouse gastruloid development between this study (left) and Stelloo et al^45^. (right).

**Extended Data Figure 5.**
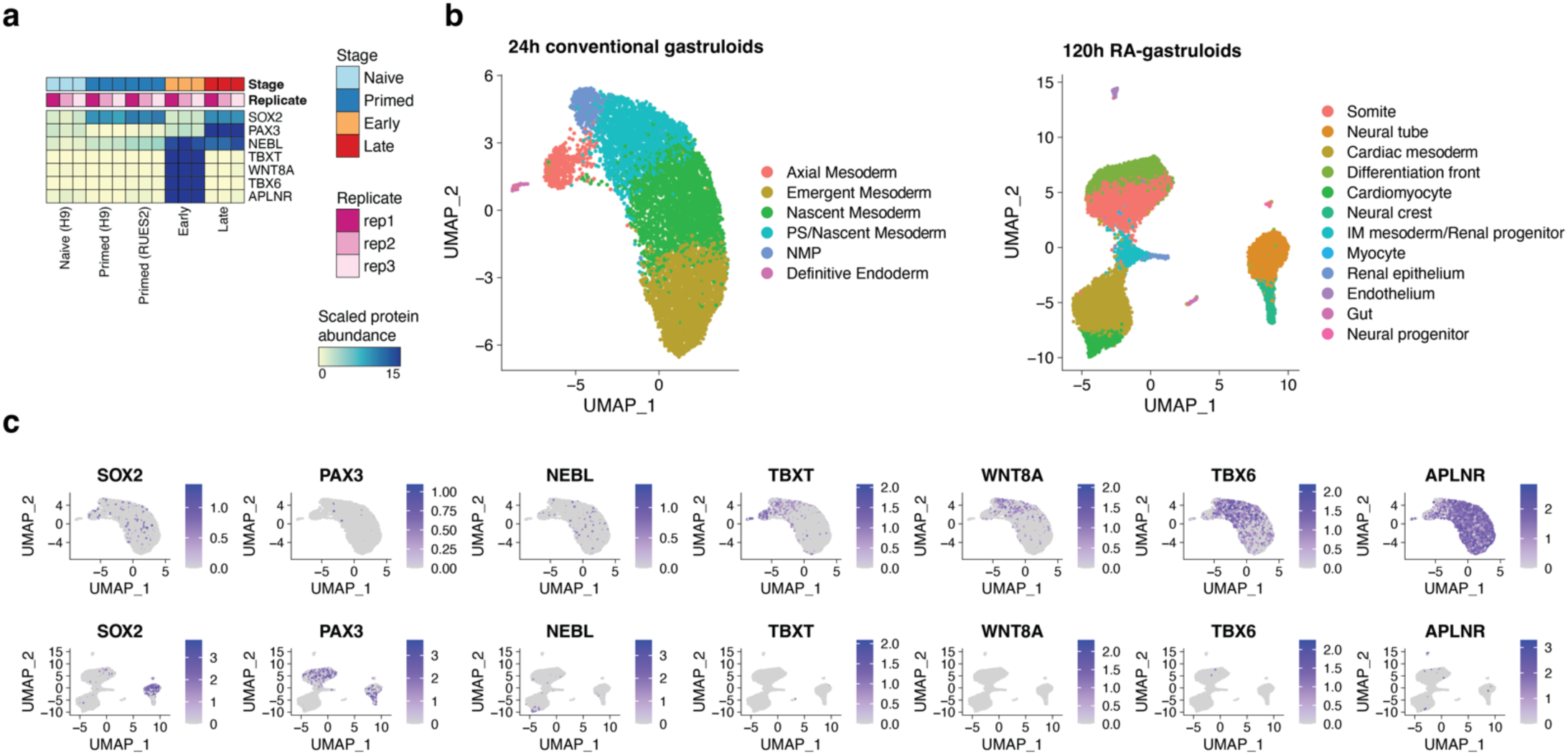
Mapping cell types contributing to bulk proteomic observations. **(a)** Heatmap depicting the temporal profiles of SOX2, PAX3, NEBL, TBXT, WNT8A, TBX6, and APLNR. Color scale for protein data indicates scaled TMT abundance. **(b)** UMAP projection of scRNA-seq profiles from 24-hour (Early) and 120-hour (Late) time points of human gastruloids. Colors in each cell type indicate the cell type **(c)** Normalized expression of SOX2, PAX3, NEBL, TBXT, WNT8A, TBX6, and APLNR from human gastruloids at 24 (top row) and 120 (bottom row) hours of development.

### Co-regulation analysis maps cooperative protein associations to known protein complexes and pathways

The vast majority of proteins quantified here do not map to known protein complexes nor are they assigned specific biological functions during early human development. Given our observations that proteins assigned to molecular modules (e.g., oxidative phosphorylation) are coherently regulated across gastruloid development (**Fig. 2d**), we sought to explore such co-regulation at a more granular level, *e.g.* among members of a specific signaling pathway or protein complex. Functional proteomics has emerged as a powerful method to annotate and assign roles to proteins in understudied contexts^46,47^. Co-regulation analysis, based on calculating correlations of protein abundances in a pairwise fashion across experimental samples, can elucidate coordinated protein functions such as macromolecular complexes and biochemical pathways^48–52^. Correlated and anticorrelated edges within the resulting networks can reveal co-regulatory effects including direct protein interactions^53^, mechanisms of action for signaling networks^54,55^, and cell state-specific roles^56^.

To apply co-regulation analysis to our data, we calculated correlations (r_Pearson_) between all 19.6 million possible pairs of the 6,261 proteins that were successfully detected and quantified in all 18 primed ESC or gastruloid samples ([3 human RUES2-GLR stages + 3 mouse stages] x 3 biological replicates). For example, proteins within known complexes were highly correlated, *e.g.* TUBG1 and TUBGCP2, which constitute the Ɣ-tubulin ring complex^57^, were highly correlated, while TUBG1 was anticorrelated with the Na+/K+ transporting ATPase, ATP1A1 (**Fig. 3a**). Across all pairs, we observed a bimodal distribution of r_Pearson_, while a similar analysis after permuting sample IDs for individual proteins yielded a normal distribution of r_Pearson_ centered at zero (**Fig. 3a**).

We focused on pairs that were either strongly correlated (R >=0.95) or anticorrelated (R <= -0.95) at a false discovery rate (FDR) of 1% (**Fig. 3c**). The resulting network consisted of 5,681 nodes (proteins) and 489,417 significant correlations or edges, of which 62% were positively and 38% were negatively correlated (174±195 edges per protein; **Fig. 3c**; **Extended Data Fig. 6a,b**; **Supplementary Table 3**). We subsetted our network to 5,227 proteins by retaining only the canonical Uniprot protein isoforms detected in our datasets and validated positively correlated edges by mapping the resulting network onto the databases cataloging known gene ontologies^42^, subcellular localizations^30^, biochemical pathways^58–62^, protein-protein interactions^63^ and protein complexes^64,65^. The proportion of annotated edges that were positively correlated edges varied by database, *e.g.* 73–92% for proteins with shared GO annotations, subcellular localization or pathway databases, but 93% for proteins previously reported to interact, and 97% for proteins belonging to the same complex (**Extended Data Fig. 6c**; **Supplementary Table 4**).

Of the positively correlated edges in the trimmed network, 37.8% were explained by at least one established annotation, a 1.4-fold enrichment over the 26.7% of all possible edges involving these 5,227 proteins that are annotated in these databases (**Extended Data Fig. 6d,e**). This was consistent with previous studies that attributed 34–42% of protein correlation network edges to previous annotations. Notably, those studies also required 41–375 different cell lines to generate co-regulation networks^53,56^. However, specific categories of annotation were much more enriched than others. For example, our network’s edges were only modestly enriched for shared subcellular localization (1.5-fold), but were strongly enriched for annotated protein-protein interactions (4.5-fold) and shared membership in a protein complex (7.4-fold) (**Fig. 3e**).

Given the relatively high proportion of positively correlated edges corresponding to protein-protein interactions and macromolecular complexes, we leveraged the untrimmed network to map positively protein pairs to specific developmental genes or protein complexes (**Extended Data Fig. 6e,f**). Anecdotally, many known protein-protein interactions were recovered. For example, BMP1, the metalloprotease that plays roles in the formation of the extracellular matrix including the processing of procollagens to their active fibril forms^66^, was highly correlated with collagens COL1A1 and COL1A2, while RPL7A, a constituent of the large ribosomal subunit, was highly correlated with other members of the large ribosomal subunit as well as tRNA synthetases (AARS1, TARS1, YARS1) that charge tRNAs with their cognate amino acids prior to translation (**Fig. 3e**).

To systematically ask whether the correlation network recovered known protein complexes, we focused on 1,357 complexes from CORUM^64^ or ComplexPortal^65^ with 3+ subunits represented in our correlation network. For this subset, an average of 80% of complex members were represented among the 5,681 proteins in the network (**Extended Data Fig. 6g**). For example, 29 of 33 (88%) of 26S proteasome complex proteins were represented, with 87% of all possible edges among these proteins detected, 100% of which were positively correlated (**Fig. 3h**). Similar trends were observed for core metabolic modules, including in the citric acid cycle, for which 90% of edges connecting pathway members were positively correlated, with only ACO1 and ACLY participating in anticorrelated edges (**Fig. 3g**).

In addition to recovering previously supported protein-protein relationships (37.8% of filtered network, **Fig. 3e**), we also nominated potentially novel relationships. Many of these novel edges are potentially driven by aspects of cell state that are unique to gastruloids and early development relative to the steady states of the workhorse cell lines that are the primary source material for the databases to which we compared our network^53,56^. Drawing from previous high-throughput proteomics studies^50,63^, we defined a protein cooperativity metric to enrich for first-degree neighbors of known complexes and pathways, termed “cooperative edges” (**Extended Data Fig. 6h**). To evaluate this approach, when members of a complex were withheld from our analysis, our cooperative edge mapping framework should recover their association to the remaining protein complex network. For example, when we divided ribosomal proteins into 60S large and 40S small ribosomal subunit groups and asked which proteins were cooperatively associated with the 40S small ribosomal subunit, we find that among the top 5 most significant hits are the 60S complex members RPL5, RPL13A and RPL32 (**Extended Data Fig. 6h**).

With this framework, we identified 1,385 cooperative proteins associating with 218 ComplexPortal complexes^65^ and 1,944 cooperative proteins associating with 524 CORUM complexes^64^ (**Supplementary Table 5**). The number of cooperative proteins per complex varied widely and was not correlated with the number of complex subunits (**Fig. 3h-k**; **Extended Data Fig. 6i**). The number of complexes that a given protein is cooperatively associated with also varied widely (**Extended Data Fig. 6j**). When comparing cooperative protein-complex relationships with protein-protein interaction databases including BioGrid and BioPlex network of interactors^63^, we found that 1,610 cooperative edges (involving 18.5% of cooperative proteins) were annotated as direct physical interactions (**Extended Data Fig. 6k**). An illustrative example involves the Chaperonin-containing T (TRiC/CCT) complex, for which 5 (13%) of the 36 most significantly cooperative proteins were primary BioPlex interactors, and 9 (25%) were BioGrid interactors (**Fig. 3i,j**). In summary, cooperative protein analysis recovered both known physical interactions as well as potentially novel associations between complexes and cooperative proteins.

We reasoned that overlaps of cooperative proteins shared by multiple complexes might inform these proteins’ functional roles. To quantify such sharing, we calculated Jaccard similarity coefficients between pairs of complexes—a perfect overlap of cooperative proteins between a given pair of complexes would result in a Jaccard similarity of 1 (**Fig. 3i**). Although most pairwise comparisons yielded little to no overlap in cooperative proteins, those that did were highly structured (**Fig. 3m**). For example, exosome complexes and histone acetyltransferase complexes each largely exhibited discrete sets of cooperative proteins that overlapped with one another but not with other complexes. The 40S and 60S ribosomal subunits, while sharing extensive overlap in terms of their cooperative proteins, also shared overlaps with the 26S proteasome and the TRiC/CCT complex (**Fig. 3m**; **Supplementary Table 6**). Other subsets of protein complexes exhibited varying degrees of sharing. For example, all SWI/SNF complexes shared cooperative proteins with other SWI/SNF complexes, but a subset of these also shared cooperative proteins with tethering complexes, as well as the ATAC (Ada-two-A-containing) coactivator^67^ and histone methyltransferase complexes (**Fig. 3m**).

**Extended Data Figure 6.**
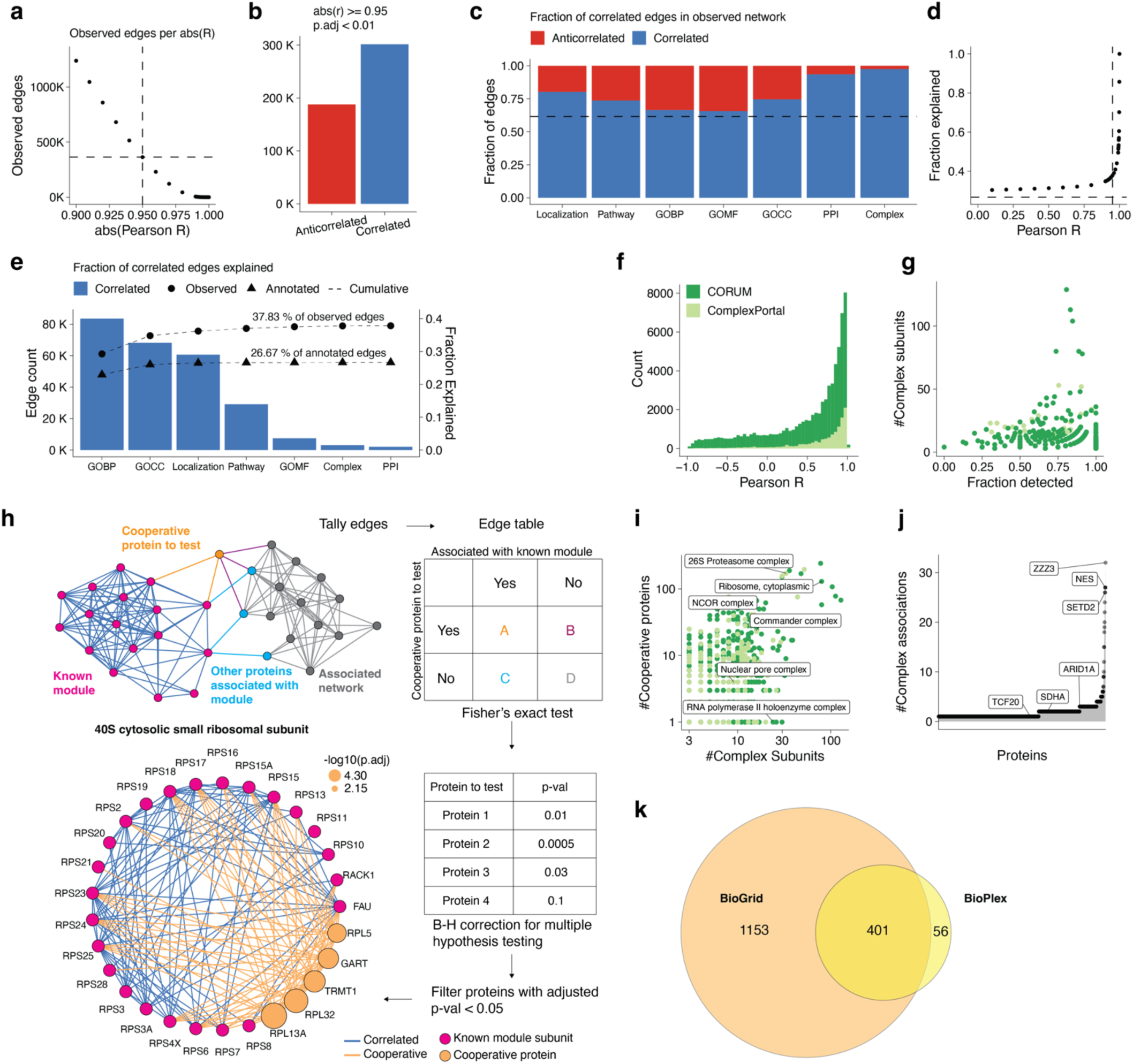
Mapping pairwise protein co-regulation onto known protein modules identifies cooperative protein associations across gastruloid development. **(a)** Number of observed edges (y-axis) in the correlation network as a function of absolute r_Pearson_ (x-axis). **(b)** Summary of correlated and anticorrelated edges in the network. **(c)** Fraction of correlated and anticorrelated edges stratified by GOBP, GOCC, Localization, Pathway, GOMF, BioPlex PPI, and Complex databases. Dashed line indicates the fraction of positively correlated edges in the trimmed network. **(d)** Fraction of correlated edges in the trimmed network explained by at least one database (y-axis) as a function of r_Pearson_ (x-axis). Horizontal and vertical dashed lines respectively indicate the fraction of edges explained in annotated network and r_Pearson_ at 0.95. **(e)** Summary of correlated edges in the trimmed network explained by shared membership in a Gene Ontology biological process (GOBP), cellular component (GOCC), molecular function (GOMF), localization, pathway, protein-protein interaction (BioPlex) or protein complex. Dotted line indicates the cumulative number of edges explained within the observed (circles) and annotated (triangle) networks.**(f)** Distribution of r_Pearson_ for protein pairs in CORUM and ComplexPortal complexes. **(g)** Fraction of complexes detected in correlation network (x-axis) versus size of protein complex (y-axis). Dots colored by database used to curate the protein complexes. **(h)** Workflow to map cooperative proteins associated with detected modules. **(i)** Number of cooperative proteins detected (y-axis) as a function of complex size (x-axis). **(j)** The number of annotated ComplexPortal complexes that were found to be cooperative with each individual protein in the correlation analysis (x-axis), *e.g.* ZZZ3 was assigned as a cooperative protein to 32 ComplexPortal protein complexes. **(k)** Venn diagram indicating the number of cooperative proteins with physical protein-protein interaction evidence to at least one subunit of their associated complex.

### Gastruloid stages and gene modules exhibit varying degrees of RNA-protein discordance

Previous studies spanning various biological contexts have reported varying extents of concordance between mRNA and protein levels^15,16,18,53,68^. Using the matched bulk RNA-seq data we acquired for these same samples, we next sought to assess the extent to which transcript abundances were predictive of protein levels in developing gastruloids. As noted above, our transcriptome data matched expectation with respect to temporal trends and stage-specific markers (**Fig. 1d**). Of note, HOX genes^69^ turned on with gastruloid induction in both species, both at the early stage in human gastruloids and the late stage in mouse gastruloids (**Extended Data Fig. 7a**).

We next calculated RNA-protein correlations of individual genes. Across the 6,010 genes with both protein and RNA data for all sampled timepoints in both species, Pearson correlation coefficients were biased towards positive correlation, consistent with previous work^53^ (mean r_Pearson_ = 0.39; **Fig. 4a**; **Supplementary Table 7**). When highly correlated (r_Pearson_ >=0.75) or anticorrelated (r_Pearson_ <=-0.75), RNA-protein relationships were stratified by broad gene classes^70–73^, *e.g.* genes associated with transcription (*e.g*. SOX2) tended to be positively correlated while those associated with the ribosome (*e.g.* NSA2) tended to be anticorrelated (**Extended Data Fig. 7b,c**). At the level of Gene Ontology (GO) biological processes, genes exhibiting highly positive RNA-protein correlation in our dataset were enriched for cytoskeletal and organ morphogenesis terms, suggesting that RNA levels are a reasonable proxy for protein abundance for these processes (**Fig. 4b**; **Supplementary Table 7**). Drilling down further to the level of complex and pathways, complexes involved in transcription (*e.g.* SOX2-OCT4 complex, CTNNB1-EPCAM-FHL2-LEF1 complex and the mRNA decapping complex) and signaling pathways (WNT and MAPK signaling) tended to be positively correlated (**Fig. 4d,e**).

**Figure 4.**
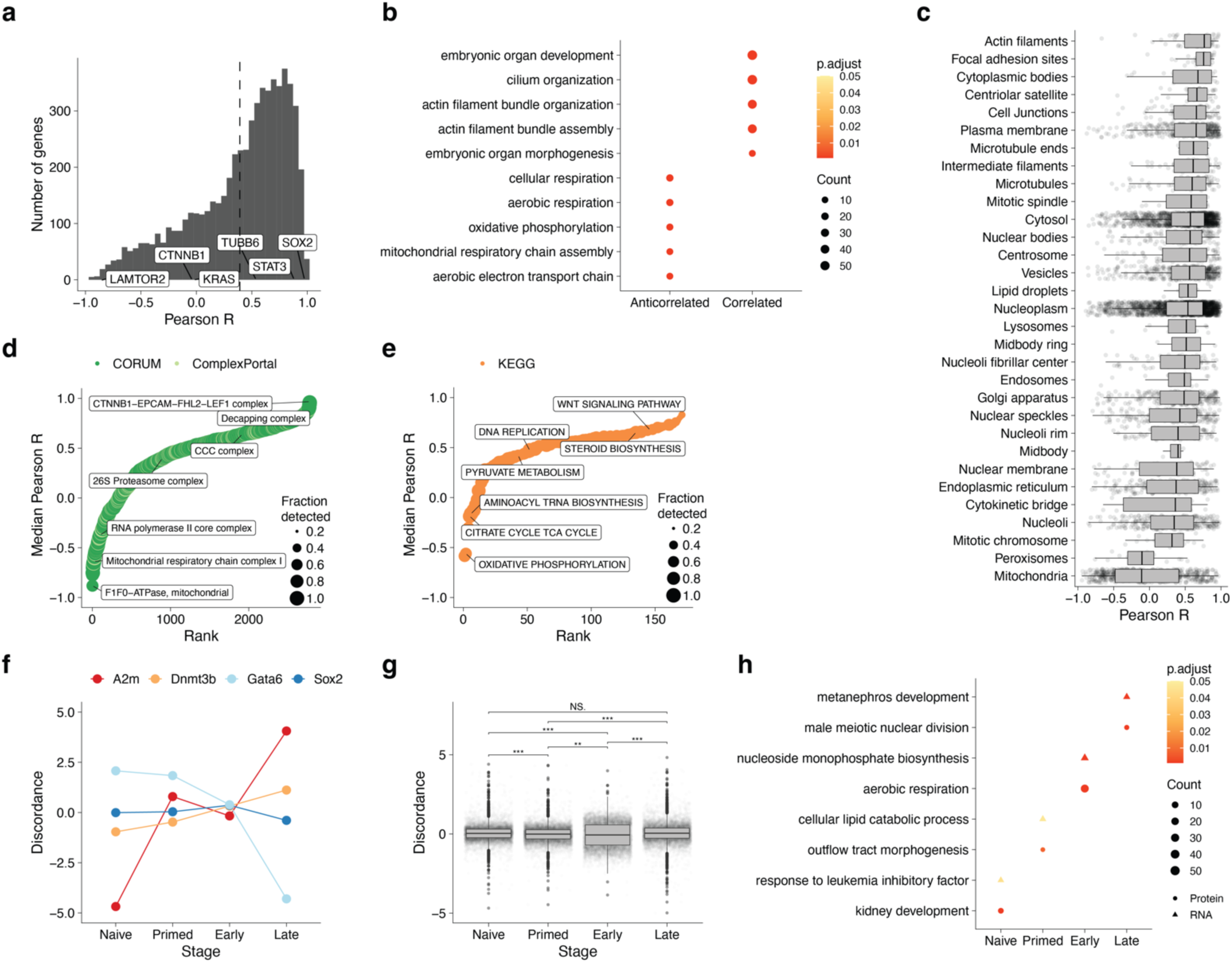
Gastruloid stages and gene modules exhibit varying degrees of RNA-protein discordance. **(a)** Histogram of correlations (r_Pearson_) between protein and RNA expression for all genes detected at the transcript and protein level in our samplings of human and mouse gastruloid development. The dashed line indicates the mean r_Pearson_ across all genes. Representative genes with varying extents of correlation are highlighted. **(b)** GO term dotplot highlighting GO-defined biological processes exhibiting high RNA-protein correlation (r >= 0.75) or anticorrelation (r <= -0.75). **(c)** Boxplot depicting the distribution of protein-RNA correlation (x-axis) as a function of subcellular location (y-axis). Rank plots of median RNA-protein r_Pearson_ across **(d)** protein complexes or **(e)** biochemical pathways. Colors indicate databases from which the module sets were curated. **(f)** Representative examples of RNA-protein discordance profiles (for any given gene, mean across replicate is shown) for various stages. **(g)** Boxplot depicting the distributions of RNA-protein discordances (for any given gene, mean across replicates is shown) for various mouse stages. **(h)** Dotplot highlighting the biological processes significantly enriched in genes exhibiting protein-abundant (circles; discordance >=1) or RNA-abundant (triangles; discordance <=-1) RNA-protein discordance. Color scale indicates the p-value adjusted using the Benjamini-Hochberg procedure and sizes of dots indicate the number of genes detected within each term.

On the other hand, at the level of GO biological processes as well as shared subcellular localization (Human Protein Atlas^30^), mitochondrial genes, particularly those involved in the electron transport chain and oxidative phosphorylation, tended to have anticorrelated RNA and protein levels (**Fig. 4b,c**), consistent with post-transcriptional and post-translational control of mitochondrial protein levels during development^74,75^. Once again drilling down further, this trend was driven by mitochondrial protein complexes (*e.g.* F1-F0 ATPase and complex I) and pathways involved in central metabolism (*e.g.* TCA cycle and oxidative phosphorylation) (**Fig. 4d,e**; **Supplementary Table 8**). In the case of Complex I, previous work in HeLa cells^76^ demonstrated that proteins in this complex were rapidly degraded post-translationally, suggesting that these systems are regulated in a similar fashion during gastruloid development.

We next sought to better understand the relationship between RNA and protein abundance as a function of developmental stage. Across all genes within each stage, we found that early mouse gastruloids exhibited substantially lower RNA-protein correlation than all other human or mouse stages (r_Pearson_ = 0.26; **Extended Data Fig. 7e**). We defined a metric of discordance between RNA and protein measurements—the log2 transformed ratio of the average fold change of a protein to its corresponding RNA—at a given stage of gastruloid development. Thus, discordance values close to 0 signify comparable levels of RNA and protein, positive discordance implies that the protein is more abundant than its corresponding transcript and vice versa. Focusing on mouse gastruloids (where for our data, all stages arose from the same cell line), Gata6 discordance was high at the naïve ESC stage (higher than expected protein, given RNA levels); while in late gastruloids, Gata6 protein-RNA discordance was low (**Fig. 4f**). In contrast, SOX2 transcript and protein abundance remained relatively consistent over time (**Fig. 4f**).

Overall, we observed varying profiles of discordance across mouse gastruloid development (**Fig. 4g, Extended Data Fig. 7e**). Applying GO enrichment analysis to genes with absolute discordance ratios greater than 1 (*i.e.* protein either highly more or less abundant than expected, given RNA levels), we observed that mitochondrial and metabolic processes tended to be discordant mainly in early mouse gastruloids (**Fig. 4h**; **Supplementary Table 9**). To understand discordance at the gene module level, we calculated the median RNA-protein discordance among genes belonging to a particular protein complex. The distributions across the four developmental states were centered at 0 (**Fig. Extended Data Fig. 7g**). We next compared the fold changes of RNA and proteins between two temporally adjacent stages with the aim of delineating when discordance emerges or resolves (**Extended Data Fig. 7h**). We found that most complexes had no significant differences in discordance between stages (*e.g.* core Mediator complex, **Extended Data Fig. 6h**). However, a minority did, *e.g.* 12% (33/279) of the protein complexes analyzed exhibited significantly different RNA and protein fold changes when comparing early vs. late stages of mouse gastruloid development. These included cytoplasmic and mitochondrial ribosomal subunits, intraflagellar transport complex B, and Complex I of the oxidative phosphorylation pathway (**Extended Data Fig. 7f,h**).

Finally, we sought to assess whether the protein levels of developmental transcription factors (TFs) could be used to adjudicate potential targets (**Extended Data Fig. 8a**). For this, we focused on Sox2, Sox3, Tfap2c, and Gata6, which exhibit distinct patterns of stage-specific protein expression during mouse gastruloid differentiation (**Extended Data Fig. 8b**). Anecdotally, transcripts for established targets of each of these TFs were indeed upregulated in a corresponding pattern, *e.g. Nanog* with Sox2, *Top2a* with Sox3, *Dppa3* with Tfap2c, and *Sox17* with Gata6 (**Extended Data Fig. 8c**)^77–82^. Although each of these TF has thousands of targets according to databases such as TFlink^83^, the RNA levels of only a subset of these are well-correlated with the TF’s protein levels in our data (r_Pearson_ >= 0.9), *e.g.* 582 for Sox2 (3.4% of its targets), 122 for Sox3 (2.6% of its targets), 218 for Tfap2c (1.5% of its targets), and 347 targets for Gata6 (6.6% of its targets) (**Extended Data Fig. 8d**). Gata6 targets were enriched for biological processes associated with SMAD signal transduction, heart development and embryonic morphogenesis, Sox2 targets were enriched for lysosome organization, autophagy, and response to LIF, while Sox3 targets were enriched for mitochondrial translation, and RNA processing (**Extended Data Fig. 8e**). Given that Sox2 exhibited elevated levels in Naive and Early stages, we asked how discrete these sets were and if the same downstream targets were upregulated at both stages. We found that out of the 245 Naive Sox2 targets and 298 Early Sox2 targets, 69 were enriched in both stages (**Extended Data Fig. 8f**). GO analyses on the discrete Sox2 targets revealed that Naive targets were enriched for response to LIF, while Early targets were enriched for processes associated with cell adhesion, placenta development and meiosis (**Extended Data Fig. 8g**). We found that downstream targets of these 4 TFs were also enriched for protein-protein interactions (**Extended Data Fig. 8h,i**). Thus, the temporal relationship between TFs and transcripts suggests that among the large number of putative targets^83^, these subsets of transcripts would be good candidates for additional study in the context of differentiating gastruloids.

**Extended Data Figure 7.**
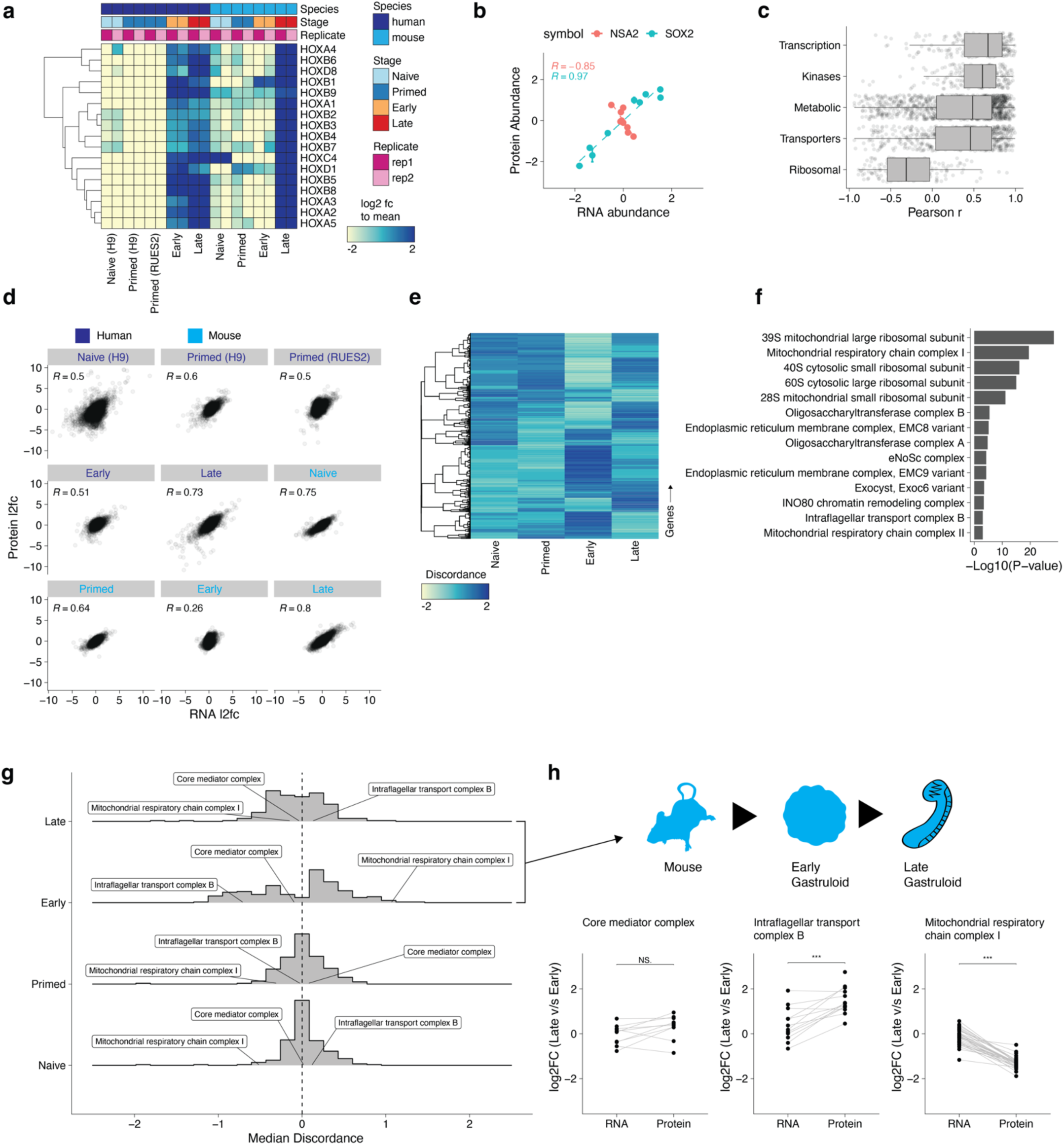
Identifying patterns of RNA-protein discordance across gastruloid development. **(a)** The temporal dynamics of the HOX gene expression cluster. Rows indicate genes, while columns signify samples. Color scale represents the log2 fold change of transcripts normalized to each sample’s respective species mean. **(b)** Representative examples of RNA vs. protein abundance correlation for SOX2 (red) and LAMTOR2 (teal). **(c)** Distributions of RNA-protein correlations (r_Pearson_) for gene sets grouped by protein class (curated from Human Protein Atlas). **(d)** Scatterplot of RNA (x-axis) and protein (y-axis) abundance across stages of mouse or human gastruloid development. **(e)** Hierarchical clustering of patterns of RNA-protein discordance ratios across genes during mouse gastruloid development. **(f)** Protein complexes whose RNA abundances differ significantly from their protein abundances when comparing early vs. late mouse gastruloids. **(g)** Median RNA-protein discordances of members of protein complexes at each stage of mouse gastruloid development. **(h)** Comparison of the RNA and protein log_2_-scaled fold-changes between early vs. late mouse gastruloids in the Mediator complex (left), intraflagellar transport complex B (middle), and mitochondrial Complex I of the oxidative phosphorylation pathway. Significance testing on RNA and protein distributions was performed using a standard t-test.

**Extended Data Figure 8.**
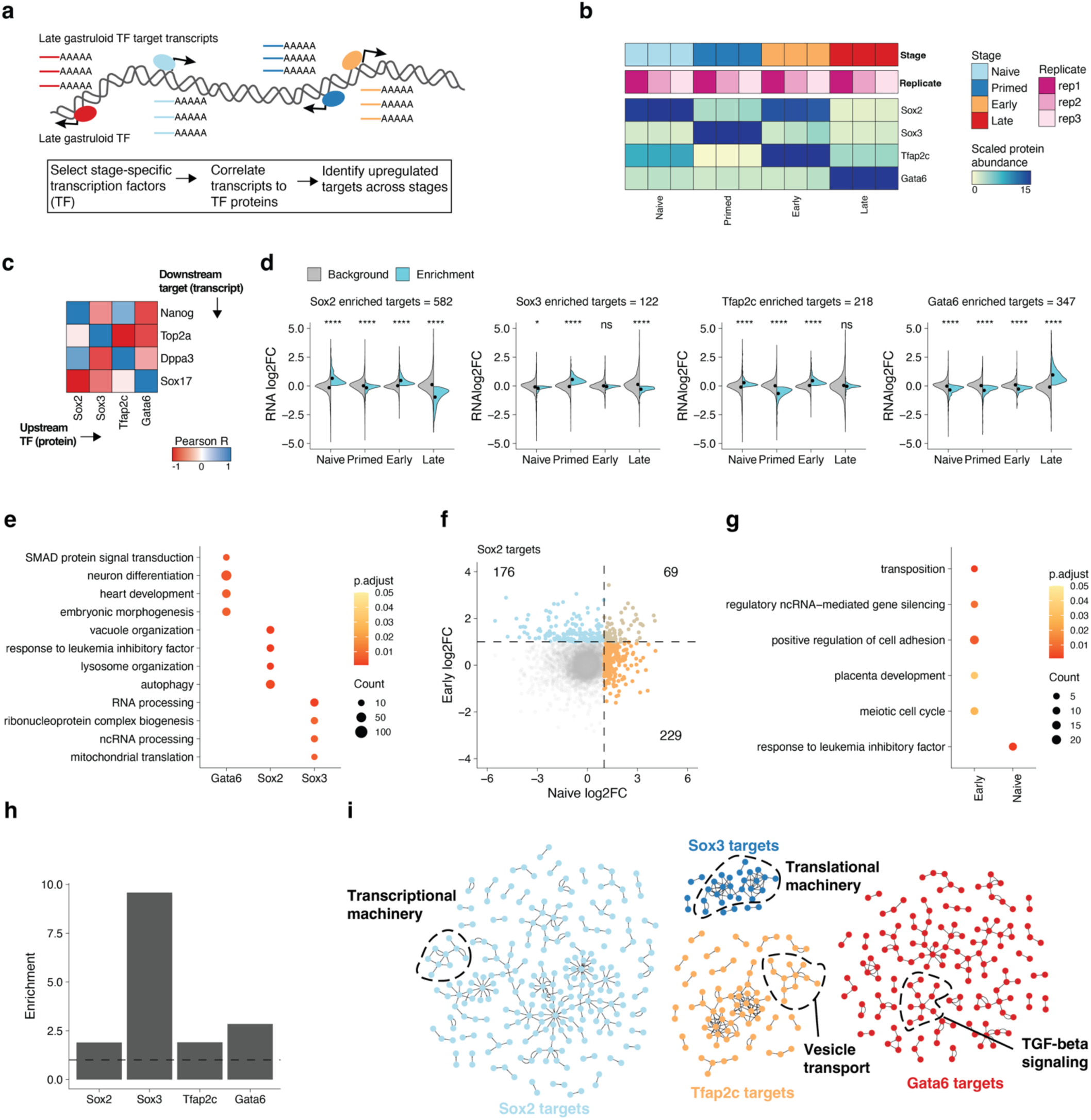
**(a)** Workflow for identifying putative downstream targets of stage-specific transcription factors. **(b)** Stage-specific protein expression of Sox2, Sox3, Tfap2c and Gata6. **(c)** Representative heatmap depicting the r_Pearson_ correlation coefficients of transcription factor protein abundance (columns) to downstream target transcripts (rows). **(d)** RNA abundance distributions (y-axes) of target transcripts to aforementioned transcription factors (top). Colors indicate the enriched (cyan) or background (gray) target transcripts to the corresponding transcription factor. Significance estimated using ANOVA (n.s. denotes not significant; * denotes p < 0.05; **** denotes p < 1.3e-8). **(e)** Dotplot highlighting the biological processes significantly enriched in downstream targets of Gata6, Sox2, and Sox3. Color scale indicates the p-value adjusted using the Benjamini-Hochberg procedure and sizes of dots indicate the number of genes detected within each term. **(f)** Scatterplot comparing levels of downstream Sox2 targets in Naive and Early gastruloid stages. Naive and Early stage enriched targets colored in orange and blue respectively while brown points indicate enriched Sox2 targets upregulated in both Naive and Early stages. **(g)** Dotplot highlighting the biological processes significantly enriched in downstream Naive and Early targets of Sox2. Color scale indicates the p-value adjusted using the Benjamini-Hochberg procedure and sizes of dots indicate the number of genes detected within each term. **(h)** Enrichment of BioPlex protein-protein interactions across Sox2, Sox3, Tfap2c, and Gata6. The dotted line indicates the background rate of protein-protein interactions in BioPlex. **(i)** Network representation of protein-protein interactions in the enriched targets of Sox2, Sox3, Tfap2c, and Gata6.

### Quantitative phosphoproteomics reveals kinase activities across gastruloid development

Developmental programs are largely driven by core signaling pathways that are dynamically regulated via phosphorylation^26^. We applied time-resolved quantitative phosphoproteomics to our sample set to map how post-translational states of proteins change across gastruloid development from ∼25μg of starting material per sample (**Fig. 1a,b**; **Extended Data Fig. 9a**; **Extended Data Fig. 10a,b**; **Supplementary Table 10**). Human and mouse phosphosites were correlated with their protein abundances (human: median r_Pearson_ = 0.71; mouse: median r_Pearson_ = 0.84) and included residues of known stem cell markers (**Extended Data Fig. 10c,d**). For example, phosphorylation of T35 and S207 on the transcription factor UTF1 decreased markedly through gastruloid development^39,84^(**Extended Data Fig. 8b**). Immunofluorescence confirmed that H2AX S139/S140 phosphorylation dynamically changes across human gastruloid development (**Extended Data Fig. 9e**). H2AX S139/S140 phosphorylation was highest in RUES2 primed ESCs, lower in H9 primed ESCs, and markedly reduced in early gastruloids before increasing again in late-stage gastruloids (**Extended Data Fig. 9f**). To further confirm our ability to decipher temporal dynamics of phosphosignaling, we leveraged that mouse gastruloids were treated with Chiron, a GSK3 kinase inhibitor that activates WNT^85–87^ and found that Gsk3a-activating phosphorylation at Y279 was inversely correlated with Chiron treatment, reflecting Chiron-dependent perturbation of Gsk3a activity during mouse gastruloid induction (**Extended Data Fig. 10e**). Additionally, kinase-substrate enrichment analysis^88–91^ identified reduced activity of GSK3B and DYRK2 during gastruloid development^45^ and increased inhibitory N-terminal phosphorylation of GSK3B^92–94^.

Based on the role of phosphosignaling in key developmental transcriptional programs, we mapped phosphosites on targets of pluripotency markers POU5F1, SOX2 and NANOG curated from previous studies^40,41^ (**Supplementary Table 11**). Fourteen proteins were shared targets of the pluripotency markers POU5F1, SOX2, and NANOG and had phosphorylation sites that exhibited temporal changes over the course of gastruloid development (**Extended Data Fig. 9c**). For example, compared to naïve ESCs, DPPA4 phosphorylation (T215) was more abundant in primed ESCs; however we found residues on DPYSL2 (S570 and T514) and DPYSL3 (S682 and S684) tended to have more total phosphorylation in early and late gastruloids. DPPA4 is a known marker of pluripotency^95^ while DYSL2 and DPYSL3 are associated with nervous system development^96^. TCF20, a transcriptional co-activator associated with neurodevelopmental disorders, displayed two distinct patterns with respect to its detected phosphosites. TCF20 residues S1522 and S1671 had maximal phosphosite abundance in primed ESCs, correlating with upstream pluripotency factors NANOG, POU5F1, and SOX2. However, phosphorylation of TCF20 S574, was most abundant in early and late gastruloids, when pluripotency factor abundance was low (**Extended Data Fig. 9d**).

Conserved human and mouse phosphorylation sites, including those on DYPSL2 and DNMT3B, exhibited highly consistent phosphorylation profiles across gastruloid differentiation. The neural stem cell regulator DPYSL2 T514/Dpysl2 T514 phosphosite abundances were consistent between mouse and human gastruloids. Notably, the N-terminal region of DNMT3B around the S100/S116 site has been challenging to resolve in structural studies^97^, lies outside of the methyltransferase catalytic domain of DNMT3B, and is important for DNA binding^97,98^. Contrasting these findings, HSP90AB1 S255/Hsp90ab1 S255 and ribosomal protein kinase RPS6KB1 S447/Rps6kb1 S447 displayed species-specific phosphosite dynamics during gastruloid development (**Extended Data Fig. 10f**). Kinase-substrate analysis also identified putative temporally-dependent MAPKAPK2 phosphorylation of ZFP36L1 at Ser92 and PRKCI phosphorylation of ECT2 Thr359 (**Extended Data Fig. 9g**). Protein abundance for ZFP36L1, a downstream target of NANOG, peaked in early gastruloids (**Extended Data Fig. 9d**) and ZFP36L1 Ser92 phosphorylation was correlated with MAPKAPK2’s predicted activity (**Extended Data Fig. 9f,g**). At the protein level, NANOG abundance had an inverse relationship with that of ZFP36L1. ZFP36L1 Ser92 may play a role in stabilizing ZFP36L1 levels and is associated with the degradation of pluripotency factors including NANOG^99,100^ and Ser92 phosphorylation correlated with MAPKAPK2 activity. Given this observation and ZFP36L1’s roles in embryonic development and implication in developmental defects^101^, we hypothesized that MAPKAPK2 may play functional roles in symmetry breaking and body axis formation. In the presence of the MAPKAPK2 inhibitor MK2in1 (**Extended Data Fig. 10i**), gastruloids failed to elongate and displayed multi-axis morphology with the majority of gastruloid cells expressing SOX2 at 120 hours of induction (**Fig. 7j-m**). The elevation of SOX2 levels began after 48 hours of induction (**Extended Data Fig. 10i**) and continued until the end of gastruloid induction. Thus, phosphoproteome analysis of gastruloid development, along with previous work^40,41^, identified potential routes of post-translational control of developmental chromatin regulators and transcription factors.

**Extended Data Figure 9.**
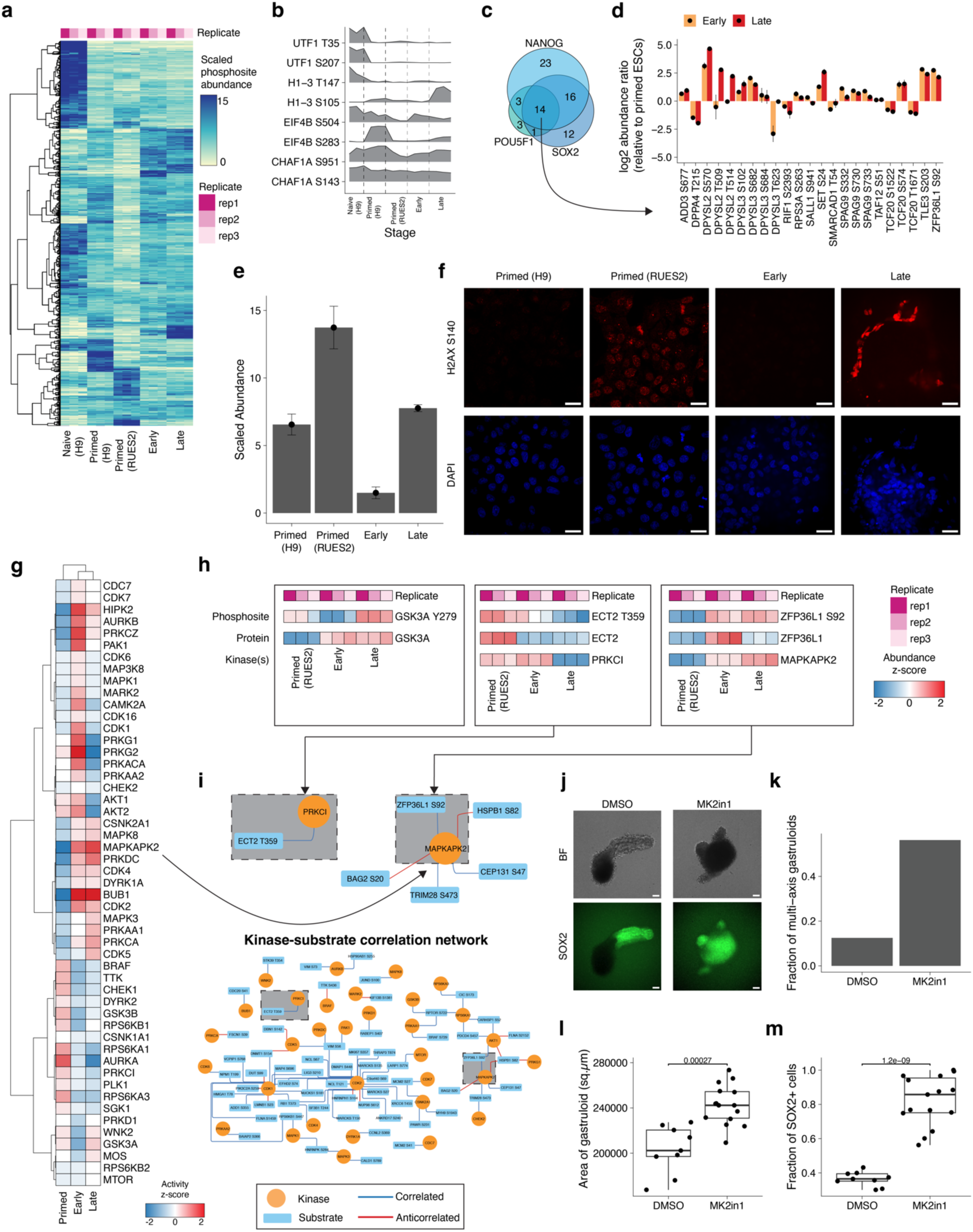
Quantitative phosphoproteomics reveals kinase activities across gastruloid development. **(a)** The temporal dynamics of phosphorylated peptides across human gastruloid development. Rows indicate phosphosites, while columns signify sample type. Color scale indicates the scaled TMT abundance of individual phosphopeptides. **(b)** Ridgeplots depicting the characteristic phosphorylation states within a given protein. **(c)** Venn diagrams depicting the detection of phosphorylated proteins that are targets of pluripotency factors SOX2, POU5F1 and NANOG. Gene sets curated from Van Hoof et al.^102^ **(d)** Phosphosites associated with downstream targets of pluripotency factors. Y-axis indicates the log_2_ abundance ratio of early (yellow) or late (red) gastruloids to primed RUES2-GLR ESCs. Mean abundance ratios are indicated with dots and error bars represent the standard deviation. **(e)** Bar plots of scaled H2AX pS139/pS140 abundance changes across human ESCs and gastruloid developmental stages. **(f)** Validation of differential phosphorylation state of pS139/pS140 (red) in Primed H9 (left) and RUES2-GLR (right) ESCs. Blue channel indicates DAPI, Scale bar: 25 µm. **(g)** Heatmap depicting the z-scores of kinase-substrate enrichment analysis. **(h)** Representative examples of temporal phosphosite dynamics in comparison to their respective proteins and cognate kinases. Color scale indicates the abundance z-score. ECT2 T359 was correlated with PRKCI, while ZFP36L1 S92 was strongly correlated with both MAPKAPK2 and AKT1. **(i)** Network of kinases (circles) connecting to their substrates (rectangles). Pairs annotated from PhosphositePlus. Edge colors indicate correlated (blue) or anticorrelated (red) relationships (absolute r_Pearson_ >= 0.5) between kinase and substrate phosphosite nodes. **(j)** Representative images of 120h gastruloids cultured with DMSO (left) and MAPKAPK2 inhibitor, MK2in1 (right). Fluorescent images depict SOX2-mCitrine expression. Scale bar: 200 µm. **(k)** Fraction of multi-axis gastruloids when treated with DMSO (control) and MK2in1 (MAPKAPK2 inhibitor). Fractions calculated from 16 gastruloid observations for each condition. **(l)** Boxplots depicting the differences in gastruloid area **(l)** and fraction of SOX2+ cells **(m)** when treated with DMSO and 10 μM MK2in1.

**Extended Data Figure 10.**
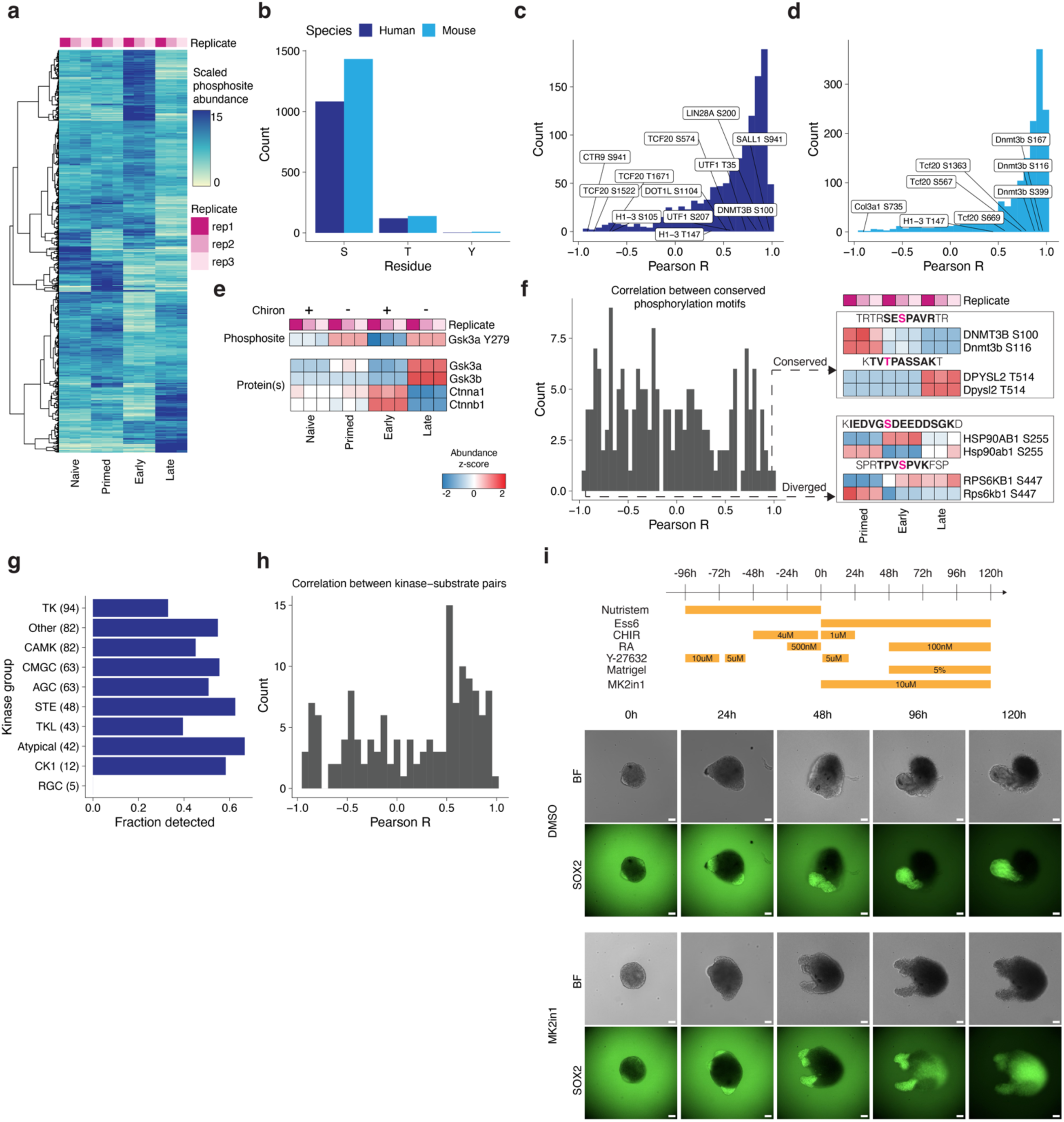
Mapping phosphorylation states across gastruloid development. **(a)** The temporal dynamics of phosphorylated peptides across mouse gastruloid development. **(b)** Number of phosphorylated sites (y-axis) identified per amino acid residue (x-axis). **(c,d)** Distribution of Pearson correlation coefficients (r_Pearson_) from comparing the abundances of phosphosites to their respective (c) human or (d) mouse proteins. **(e)** Effects of temporal Chiron treatment on protein and/or phosphorylation dynamics of Gsk3a, Gsk3b, Ctnna1, and Ctnnb1. **(f)** Distribution of r_Pearson_ computed from comparing temporal abundances of conserved phosphorylation motifs between human and mouse (left). Representative tile plots of conserved and diverged phosphosite profiles across motifs shared between humans and mice (right). Detected peptide (bold) and phosphorylated residue (magenta) are highlighted above each tile plot. **(g)** Proportion of human protein kinases detected by kinase group. Kinase annotations curated from KinMap explorer^103^. **(h)** Histogram of r_Pearson_ between human kinase-substrate pairs detected across human gastruloid development. Pairs curated from PhosphositePlus. **(i)** Protocol and timecourse of RA-gastruloid development when treated with DMSO (top row) and MAPKAPK2 inhibitor, MK2in1 (bottom row). Fluorescence images indicate the expression of SOX2-mCitrine. Ess6, Essential 6 media; CHIR9, CHIR99021; RA, Retinoic Acid. Scale bar: 200 μm.

### Co-regulatory networks of protein dynamics in gastruloids link to shared phenotypes and developmental disorders

*In vitro* models of early development offer tractable platforms to model congenital disease states and map the molecular mechanisms underlying them. To investigate the temporal dynamics of proteins linked to developmental disorders, we intersected our dataset with the Gene Curation Coalition (GenCC)^104^ and Deciphering Developmental Disorders (DDD)^105^ databases. There were 1,980 proteins (27%) quantified in our datasets with at least one disease association in at least one of these databases (**Fig. 5a**; **Supplementary Table 12**). Anecdotally, genes linked to the same disease tended to be co-regulated across gastruloid development. For example, genes associated with Leigh Syndrome, a congenital early-onset neurological disorder associated with mitochondrial dysfunction, tended to be upregulated in primed ESCs, while genes linked with broad intellectual disability mostly showed increased abundance during the gastruloid stages (**Fig. 5a**). To ask if co-regulation was a general trend among genes associated with the same disease, we calculated the mean Pearson correlation coefficient across all pairs of detected proteins associated with a given disease by GenCC, and found these tended to be positively correlated (average r_Pearson_ = 0.46, **Fig. 6b**).

**Figure 5.**
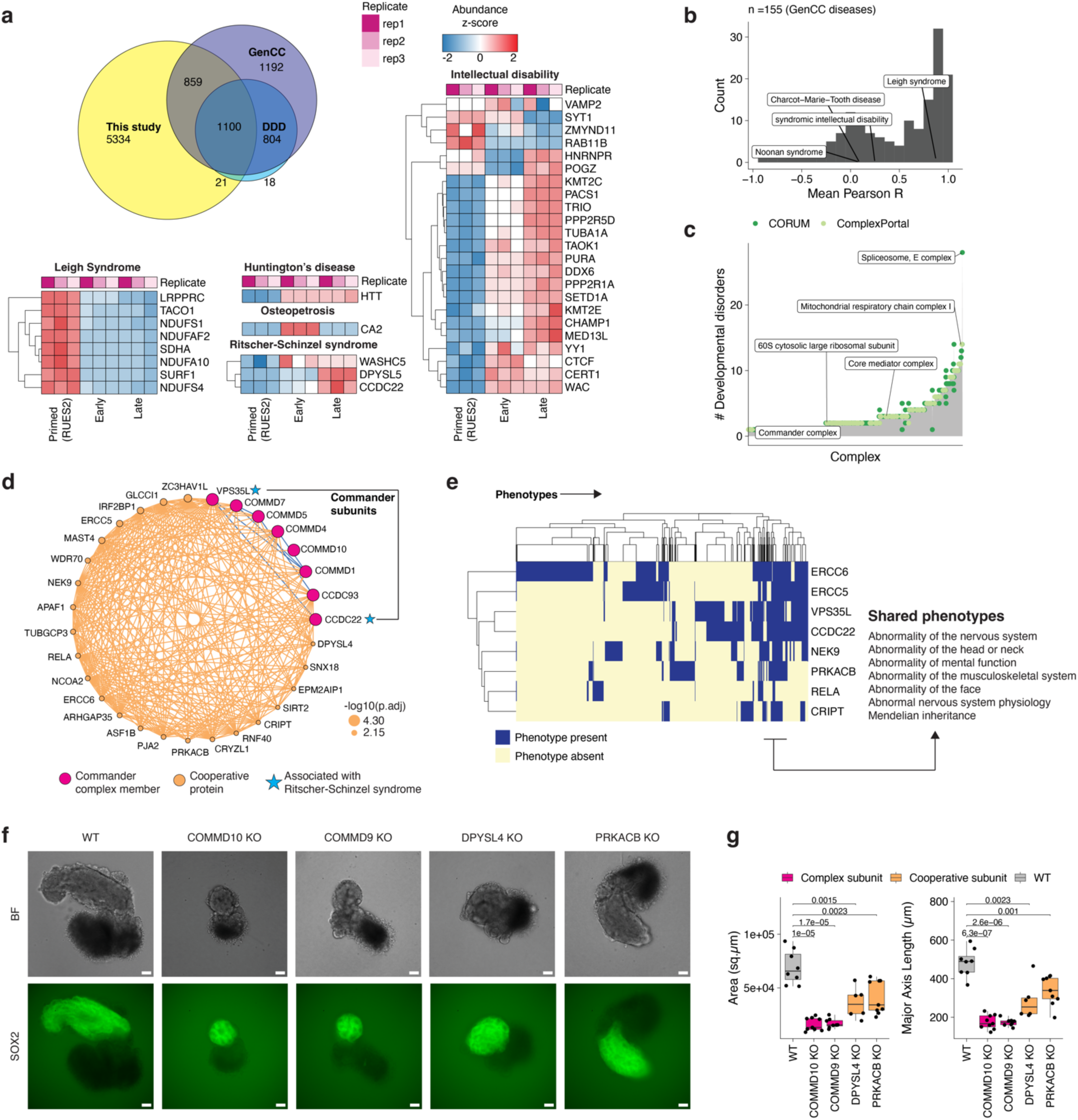
Co-regulatory networks of protein dynamics in gastruloids link to shared phenotypes and developmental disorders. **(a)** Overlap of proteins detected across our dataset, GenCC and DDD. Heatmaps correspond to the temporal abundance changes of human proteins (rows) associated with a specific developmental disorder across human gastruloid development (columns). **(b)** Distribution of mean Pearson correlation coefficients across GenCC disease sets. Only diseases with >=2 genes are plotted. Mean r_Pearson_ was calculated by averaging Pearson correlation coefficients detected pairs of proteins in our dataset. **(c)** Histogram of the number of developmental disorders associated (y-axis) with genes comprising protein complexes (x-axis). **(d)** Co-regulation network of the Commander complex subunits. Size of orange nodes indicates the significance of cooperative association (-log_10_ of the adjusted p-value; Fisher’s exact test, see Methods). Proteins associated with developmental disorders (blue stars) were linked to the Commander complex co-regulation network. **(e)** Heatmap depicting the extent of shared phenotypic overlap (columns) across genes (rows) in the Commander subnetwork. **(f)** Representative images of WT (wild-type), COMMD10 KO, COMMD9 KO, DPYSL4 KO, PRKACB KO gastruloids. n= 48 gastruloids per genotype, Scale bar: 200 μm **(g)** Boxplots comparing the area and major axis length of wild-type with genetically perturbed gastruloids. Significance determined using standard t-test.

Mapping disease-associated genes onto known protein complexes can inform their molecular roles in developmental disorders. There are 461 developmental disease-associated genes whose protein products contribute to 217 ComplexPortal and 631 CORUM complexes (**Supplementary Table 12**). Complexes were associated with an average of 2.95 ± 2.68 developmental diseases, with the spliceosome E complex and mitochondrial respiratory Complex I are associated with the largest number of developmental disorders (**Fig. 5b**). With the goal of identifying additional disease genes that might be related to these protein complexes, we leveraged our data together with the aforedescribed co-regulatory analysis heuristic (**Fig. 3a**; **Extended Data Fig. 6j**). Towards supporting these protein-complex associations, we mined BioPlex and BioGrid, which identified 232 edges linking cooperative disease proteins to CORUM complexes, and 180 edges linking cooperative disease proteins to ComplexPortal complexes (**Extended Data Fig. 11a**).

Functional proteomics can be a powerful approach to nominate molecular functions for disease-associated genes with molecular functions, as well as to advance our mechanistic understanding of how their aberrant function in specific developmental contexts might give rise to specific disease phenotypes^106,107^. To illustrate how the data reported here might be useful in this regard, we highlight examples involving Leigh Syndrome and Ritscher-Schinzel Syndrome.

Leigh syndrome is an early-onset mitochondrial neurometabolic disorder impacting the central nervous system. Its symptoms include, ataxia, developmental delay and hypotonia^108^. We found that the protein levels of 51 Leigh Syndrome-associated genes detected in our data were highly correlated with one another in our data (**Fig. 5b**, mean r_Pearson_ = 0.87). In our co-regulation network, Leigh syndrome proteins clustered with genes associated with central metabolism (*e.g*. Complex I and mitochondrial ATP synthase) and were significantly enriched in an oxidative phosphorylation co-regulation subnetwork (p < 9.6×10^-15^, Fisher’s Exact test, **Extended Data Fig. 11b**).

Ritscher-Schinzel syndrome is a developmental disorder characterized by abnormal craniofacial, cerebellar, and cardiovascular malformations, classically associated with WASHC5 and CCDC22 but more recently with VPS35L and DPYSL5 being implicated as well^109–114^. These four proteins were positively correlated in our data (mean r_Pearson_= 0.78) with two of them (CCDC22 and VPS35L) clustering within a co-regulation network involving subunits of the Commander complex (**Fig. 5c**). The Commander complex consists of two subcomplexes, CCC and Retriever^115^, and is mainly involved in trafficking of cargo including endosomal recycling of proteins^116^. Perturbations of CCC subunits COMMD9 and COMMD10 in mice have been previously linked to severe developmental defects and embryonic lethality^117,118^. While we detected all 16 Commander subunits, including the three heterotrimeric components of the Retriever complex (**Extended Data Fig. 11c**), our co-regulation network contained 7 CCC subunits, 1 Retriever subunit, and 23 cooperative proteins (**Fig. 5d**).

While we detected 2 of the 4 genes associated with Ritscher-Schinzel syndrome, we also observed 7 of the 31 proteins in the Commander coregulatory network had GenCC disease associations. We hypothesized that cooperative disease-associated proteins in the Commander network would share similar phenotypic features. To measure the extent of phenotypic overlap among disease-associated proteins in the Commander network, we leveraged the Monarch database of gene-phenotype relationships^119^. Eight proteins in the Commander coregulatory network had phenotype associations, which clustered into groups based on shared sub-phenotypes (**Fig. 5f**). For example, and unsurprisingly given how the syndrome is defined, Ritscher-Schinzel syndrome genes CCDC22 and VPS35L shared highly similar phenotypes. More broadly however, Commander coregulatory proteins exhibited overlapping phenotypic characteristics, including abnormality of the nervous system, mental function, and musculoskeletal system (**Fig. 5f**). We hypothesized that the Commander co-regulatory network could identify genes associated with developmental disorders. To test this, we perturbed 2 commander subunits (COMMD9 and COMMD10) and 2 co-regulatory proteins (DPYSL4 and PRKACB) in human ESCs and generated gastruloids from these knock-out lines. Knockout of both Commander subunits, COMMD9 and COMMD10, led to drastic morphologies with gastruloids generated from these knockout lines failing to elongate resulting in abnormal neural tube morphology (**Fig. 5f**). Perturbation of DPYSL4 resulted in similar abnormal neural tube morphologies phenocopying the knockouts of the Commander subunits (**Fig. 5f**). Perturbed gastruloids had reduced areas and a pronounced reduction in the length of major axes across the knockouts (**Fig. 5g, Extended Data Fig. 11b**). Perturbation of PRKACB also resulted in gastruloids with reduced area, though the altered major axis length was less pronounced.

**Extended Data Figure 11.**
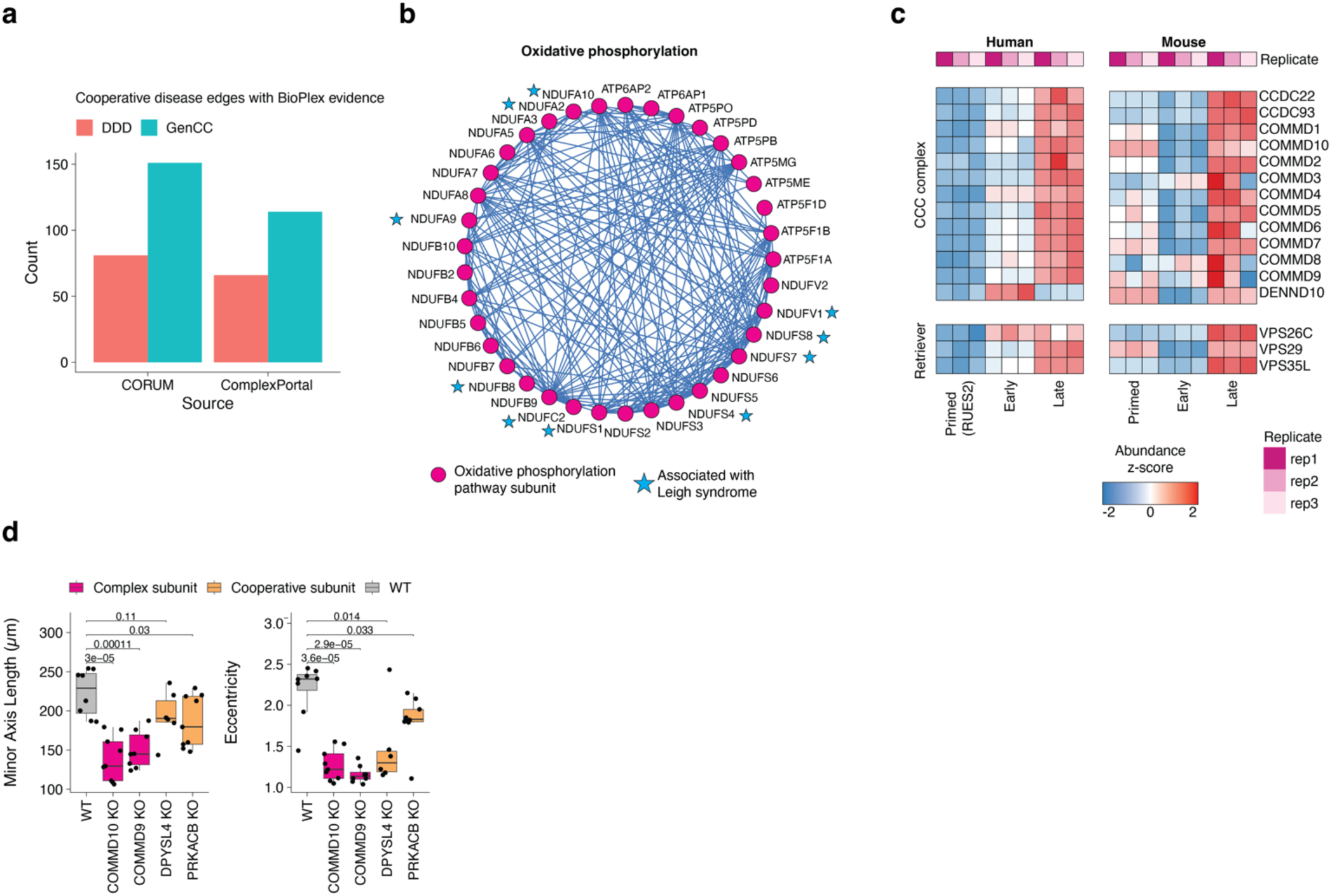
**(a)** Number of cooperative disease proteins with physical evidence in BioPlex or BioGrid to known protein complexes**. (b)** Oxidative phosphorylation co-regulatory network (pathway associations curated from WikiPathways). Proteins associated with Leigh syndrome (blue stars) were enriched in the oxidative phosphorylation co-regulation network (Pathway curated from WikiPathways). **(c)** Temporal protein profiles of the Commander complex across human and mouse gastruloid development. **(d)** Boxplots comparing the distributions of minor axis length, eccentricity (left) and major-to-minor axis ratio (Eccentricity, right) of wild-type and perturbed gastruloids. Significance determined using standard t-test.

## DISCUSSION

In this study, we leveraged the tractable and scalable properties of stem cell derived human and mouse gastruloids to systematically profile temporal changes across four key stages of their development. While the numbers of *in vitro* models of embryogenesis continue to expand and are increasingly characterized with scRNA-seq or scATAC-seq, only recently have they been subjected to phenotyping at the protein level. For example, a recent study applied mass spectrometry to map the temporal protein dynamics across stages of mouse gastruloid development, yielding insights into germ layer proteomes and associated phosphorylation states^45^. However, this study was restricted to mice and stages surrounding gastruloid induction. Thus, we sought to extend the application of these approaches to a human model of gastrulation to enable multi-species comparisons and generate a resource to explore additional developmental states mapping to pre- and post-implantation to gain a more comprehensive view of mammalian gastrulation.

We generated an integrated proteomic, transcriptomic, and phosphoproteomic dataset in both mouse and human gastruloids to define the molecular dynamics of this model of early development. At a metabolic level, we observed over human gastruloid development, that proteins in the TCA cycle tend to be upregulated in primed ESCs as compared to late gastruloids, and late gastruloids display elevated levels of glycolytic proteins. These results suggest a refocusing of cellular composition on metabolic energy production in support of larger organismal changes, in broad agreement with previous studies demonstrating the metabolic shift to glycolysis in post-implantation embryos^120,121^. It is important to note that while previous studies using transcriptomic approaches suggest a bivalent metabolic state in epiblast cells^121^, we found that primed RUES2-GLR cells had increased abundance of oxidative phosphorylation proteins. Additionally, RUES2-GLR proteomes when compared to primed H9 cells displayed an elevated oxidative phosphorylation profile and stem cell line-specific metabolic states. However, the downregulation of oxidative phosphorylation was observed in gastruloids generated from either REUS2-GLR or H9 cells suggesting shifts in metabolism underlie the formation of early stage gastruloids. The elevated levels of glycolysis in later stages of gastruloid development are in line with previous studies demonstrating that elevated levels of glycolysis underlie somite formation and occurring in human RA-gastruloids from 96-120hrs after induction^122–124^. Future efforts profiling in neural or somite organoid models can shed light on how metabolic states govern their differentiation.

Our data enabled comparison of temporal dynamics of protein expression and conservation (or lack thereof) across human and mouse gastruloid development. While late gastruloids across both species were only modestly correlated, key developmental genes often displayed conserved patterns of expression. For example, in both species the protein abundance for pluripotency markers POU5F1, NANOG, CDH1 were all lower in late gastruloids compared to their stem cell progenitors. Conversely, ZEB2 (a key protein involved in epithelial-mesenchymal transition^125^), SOX9 (a neural crest marker), CDX2 (a caudal axial stem cell marker), and MEIS1 (a cardiomyocyte marker) protein abundances all increased at the protein level in both human and mouse gastruloids. We found conserved upregulated biological processes include regulation of cell differentiation, organ morphogenesis, heart/muscle development, while conserved downregulated processes include amino acid metabolism and transport. Conservation of these processes is particularly striking given substantial protocol differences (for example the 10-fold difference in starting cell number) between human and mouse gastruloid induction suggesting that these processes are conserved. Recent studies have demonstrated that starting cell number significantly impacts gastruloid developmental trajectories^126,127^, raising the questions of whether observed similarities reflect conserved developmental biology or technical limitations associated with gastruloid generation. While our study cannot confidently rule out technical variation between species, it offers a starting point for further investigations into the conservation of developmental programs in early embryogenesis.

Upon comparing the stages sampled across both species, we surprisingly found that the proteomes of primed RUES2-GLR cells were closest to those of the early mouse gastruloids. While this association was largely driven by upregulation of mitochondrial proteins, it is possible that RUES2-GLR cells (thought to be between pre- and post-implantation states) may already be primed towards gastrulation at the protein level. Our results thus also highlight potential species-specific differences in staging, particularly with respect to metabolic and mitochondrial states. However, more work is needed to understand the extent of these effects and to rule out the possibility that these are due to cell line-specific differences that do not reflect *in vivo* effects. We imagine that a more continuous sampling of early differentiation, together with computational staging between species^10^ will resolve this question. When comparing protein abundances to their corresponding transcripts, we observe modest correlation (r_Pearson_ = 0.39) with a clear discordance between mitochondrial proteins and transcripts including proteins underlying key metabolic pathways such as oxidative phosphorylation being anticorrelated. However, this was not the case for other pathways such as WNT signaling, steroid biosynthesis and glycolysis. Our findings are in broad agreement with other studies mapping RNA-protein relationships in developmental contexts across a range of organisms^16,19–23^ and highlight the roles of post-transcriptional regulation in gastruloid development as well as the need to study multiple layers of biomolecular composition of organisms during development. Future studies, applying ribosome profiling^128^ and single cell multi-omics of the proteome and transcriptome^129^ may inform the translation and turnover rates of specific proteins across developmental stages.

Studies of co-regulation of protein systems (*i.e.* complexes and pathways) during gastrulation have been limited owing to the lack of tractable and scalable models of mammalian gastrulation. Here we recover hundreds of known macromolecular protein complexes and biochemical pathways and map their dynamics during gastruloid development. In mining our co-regulatory networks of protein expression, we identify thousands of new cooperative proteins associating with existing complexes and pathways suggesting possible developmental roles in gastrulation. While many proteins were discrete to the complexes they cooperated with, we also found shared sets of cooperative proteins. For example, chromatin remodelers (SWI/SNFs and BAFs) and histone methyltransferases (SIN3A/SIN3B histone methyltransferases) and acetyltransferases (HBO complexes) shared cooperative proteins. These relationships represent a resource for further exploration using biochemical assays to disentangle those proteins that directly interact from those in shared pathways.

Finally, we found that co-regulation and network analysis were able to identify disease neighborhoods. Often we observed that genes associated with the same developmental disorders were highly correlated with one other at the protein level. Subunits of the Commander complex are associated with Ritscher-Schinzel syndrome^130,131^. The cooperative network containing this complex consisted of 2 of the 4 genes strongly linked to Ritscher-Schinzel syndrome. Proteins we identified as coregulated with the Commander complex, including DPYSL4 and PRKACB, are associated with similar phenotypic characteristics in gastruloids. DPYSL4 has been associated with neural functions including neurite initiation and dendrite growth of hippocampal neurons^132,133^ while PRKACB has been associated with neural tube defects^134^ while their molecular mechanisms and temporal roles during early embryogenesis remain understudied. Our findings highlight the value of co-regulatory protein analysis in developmental models as powerful starting points to better understand the roles of these genes in gastrulation. Finally, this study offers scalable and multidimensional approaches to provide high-content phenotyping going beyond ubiquitous nucleic acid-centric assays. For example, generalizable protein-focused approaches can be extended to phenotype genetically and chemically perturbed gastruloids along with other clinically relevant stem cell models that are increasingly being used to model embryogenesis^5^.

Though this study quantitatively profiles the transcriptome, proteome, and phosphoproteome across gastruloid development, it is not complete nor comprehensive. First, while we sample 4 stages of embryonic development in gastruloids, profiling more time points with finer windows across gastruloid development would provide greater resolution and enhance our understanding of the temporal dynamics present in these samples. Secondly, though we quantified ∼7,500 human and ∼8,700 mouse proteins, which represents a substantial portion of the observable proteome^135^, additional coverage of low abundance, developmentally associated genes could be attained using targeted, quantitative mass spectrometric analyses. Thirdly, gastruloids consist of diverse cell types arising from all three germ layers. Our approaches were all bulk measurements and lacked cell-type-specific resolution. Additional characterization of separate cell types with fluorescence activated cell sorting (FACS) before phenotyping gastruloid development with proteomics^45^, could overcome this limitation. While both conserved and species-specific differences are observed between mouse and human gastruloids, the absence of standardized mammalian gastruloid culturing techniques makes it difficult to distinguish true species-specific variations in our datasets from differences that result from varying culture conditions between human and mouse gastruloids. Finally, although gastruloids are powerful surrogates to model specific characteristics of early mammalian embryogenesis, they still do not entirely reconstitute embryogenesis *in vitro*. Future multi-omic studies will lay the foundation for advancing these stem cell models to more accurately recapitulate embryogenesis and allow us to better understand the cellular and molecular mechanisms driving embryonic development.

## METHODS

### Ethics statement

All research conducted in this work including the induction, cellular and/or molecular analysis of both mouse gastruloids and human RA-gastruloids were reviewed and approved by the Embryonic Stem Cell Research Oversight of the University of Washington (E0047-001). This work was performed in compliance with the principles laid out in the International Society for Stem Cell Research Guidelines for Stem Cell Research and Clinical Applications of Stem Cells^136^. No experiments involving human embryos and gametes were performed in this study. Both human and mouse gastruloids were cultured for no longer than 5 days after induction.

### Mouse cell lines

E14Tg2a cell line was obtained from Dr. Christian Schroeter (Max Planck Institute).

### Mouse naïve ESC culture

Mouse naïve ESCs were maintained in 2iLif medium^86^ containing 3 µM CHIR99021 (Millipore-Sigma, SML1046), 1 µM PD0325901 (Stemcell Technologies, 72184), and 1,000 U/ml LIF (Millipore, ESG1107) and passaged with TrypLE (Thermo, 12604021) every other day onto new wells, which were coated with 0.01% poly-L ornithine (Millipore Sigma, P3655-10MG) and 300 ng/ml Laminin (Corning, 354232).

### Mouse EpiLC differentiation

Mouse EpiLC differentiation was performed as previously described^137^. Briefly, 1×10^5^ mouse naïve ESCs were seeded onto a well on a 12-well plate, which was coated with human plasma fibronectin (Thermo, 33016015) in EpiLC differentiation medium (N2B27 + 20 ng/ml ActivinA + 12 ng/ml bFGF + 1% KSR). The medium was changed a day after the seeding. Day2 EpiLCs were dissociated with TrypLE (Thermo, 12604021) and sampled.

### Mouse gastruloid induction

Mouse gastruloid induction was performed as previously described^9^. Briefly, mESCs cultured in 2iLiF medium were dissociated with TrypLE, and 300 cells were seeded into U-bottomed, non-adherent 96-well plates in N2B27 medium and kept for 48 hours in 37°C 5% CO_2_ incubator. After 48 hours, 150 µl of N2B27 containing 3 µM CHIR99021 was added to each well. At 72 and 96 hours, 150 µl medium was replaced with fresh N2B27 medium lacking CHIR99021. Mouse gastruloids were sampled at 72 and 144 hours after induction.

### Human cell lines

Pluripotent stem cell lines, hESCs (RUES2-GLR), were gifted by Dr. Ali Brivanlou (Rockefeller University). Chemically reset (cR) H9 naïve and primed cells were kindly gifted by Dr. Austin Smith (University of Exeter).

### Human naïve ESC culture

Chemically reset (cR) H9 naïve hESCs were propagated in N2B27 with PXGL (P-1mM PD0325901, 2mM X-XAV939, G-2mM Gö 6983 and L-10 ng/mL L-human LIF) on irradiated MEF feeders as described previously^33,138^. Y-27632 and Geltrex (0.5mL per cm^2^ surface area; Thermo Fisher Scientific, A1413302,) were added during re-plating. To remove MEF cells, cells were passaged on geltrex-coated wells with the 1 μL/cm^2^ and were repeatedly passaged by dissociation with Accutase (Biolegend, 423201) every 3-5 days for five successive passages.

### Human primed ESC culture

Human primed ESCs were cultured in StemFlex (Thermo, A3349401) on Geltrex (Thermo, A1413201) and were routinely passaged using StemPro Accutase (Thermo, A1110501) to new Geltrex-coated wells as recommended by the manufacturer. For the first 24 hrs after passaging, hESCs were cultured in StemFlex with 10 μΜ of Rho Kinase inhibitor Y-27632 (Sellek, S1049) to prevent apoptosis.

### Human RA-gastruloid induction

Human RA-gastruloids were induced as described previously^10^. Briefly, ∼2×10^4 hESCs were plated onto a single well of a Vitronectin-coated 12-well dish (Gibco, A14700) in Nutristem hPSC XF medium (Biological Industries, 05-100-1A) in the presence of 10 µM Y-27632. After 24 hours, the medium was replaced with Nutristem containing 5 µM Y-27632. At 48 hours the medium was replaced with Nutristem containing 4 µM CHIR (Millipore, SML1046). At 72 hrs, the medium was replaced with Nutristem containing 4 µM CHIR and 500 nM RA (Millipore Sigma, R2625). Pre-treated cells were detached using StemPro Accutase, dissociated into single cells suspension, and then 4,000 cells per well of a U-bottom shaped 96-well plate with 50 µl Essential 6 medium (Thermo, A1516401) containing 1 µM CHIR and 5 µM Y-27632. At 24 hrs, 150 µl of Essential 6 medium was added to each well. At 48 hrs, 150 µl of the medium was removed with a multi-channel pipette, and 150 µl of Essential 6 medium containing 5% Matrigel and 100 nM RA was added and maintained at 37°C and 5% CO_2_ until 120 hrs. Human gastruloids were sampled at 24 and 120 hours after induction.

### Perturbation experiments

#### Genetic perturbations in ESCs

Genetic perturbations in RUES2-GLR ESCs were performed as previously described using CRISPR-Cas9 RNA–protein complexes^10^. In brief, equal molar amounts of crRNA and tracrRNA (IDT, **Supplementary Table. 13**) were hybridized by heating at 95 °C for 5 min in a thermal cycler and cooling to room temperature for 10–20 min. AltR-Cas9 protein (IDT, 1081058) was added to the hybridized crRNA-tracrRNA mixture to assemble Cas9 RNPs.

RUES2-GLR ESCs were dissociated with StemPro Accutase whose activity was quenched DMEM-F12 nutrient mix supplemented with 10 mM Y-276322. For each perturbation, 200,000 cells were collected by centrifugation at 250g for 5 min. Cells were resuspended in 20 µl nucleofection buffer (16.4 µl Nucleofector Solution + 3.6 µl Supplement) provided in P3 Primary Cell 4D-Nucleofector X kit S (Lonza, V4XP-3032). 3 µl RNP and 0.5 µl of AltR-Cas9 Electroporation Enhancer (IDT, 1075915) were added to cells before transferring them into 16-well Nucleocuvette Strips and electroporated with the CA-137 nucleofection program. The nucleofected cells were transferred to a 12-well plate that contained Nutristem or StemFlex with 10 mM Y-27632 and after 24 h, the medium was replaced with Nutristem without Y-27632. Cells were maintained until they reached 50– 70% confluence. Then, the electroporated cells were transferred onto 0.5 μg/cm^2^ Vitronectin-coated 12-well plates prior to proceeding with RA-gastruloid induction steps as described above.

#### Chemical perturbations in gastruloids

10 mM stocks were prepared by resuspending MK-2-in-1 (HY-12834, MedChemExpress) in DMSO. MAPKAPK2 perturbations were performed by inducing RA-gastruloids in the presence of 10 μM MK2in1 added on day 0 and replenished on day 2.

### Immunostaining of ESCs and gastruloids

ESCs were fixed and stained as described previously^147^. Briefly, ESCs were cultured on Matrigel with StemFlex or mTeSR+ in glass-bottom 12-well plates (Cellvis, P12-1.5H-N). Cells were washed 3 times with PBS prior to a 30-minute fixation in 4% paraformaldehyde. Cells were then washed 3 times with PBS before permeabilizing with 0.1% Triton X-100 (in PBS) for 30 minutes. Cells were stained with primary antibodies diluted to the recommended working concentrations in Cell Painting Buffer^140^ (1x HBSS, 1% BSA, and 0.01% sodium azide) with 0.75% Triton-X 100 for 1 hour while shaking. Then cells were washed thrice PBST (0.2% Tween-20) and stained with secondary antibodies (diluted 1:500 or 1:1000 in Cell Painting Buffer) for 1 hour while shaking in the dark. Cells were washed thrice with PBST and kept in the dark after staining. Cells were imaged in UltraPure™ SSC (Thermo Fisher Scientific, 15557044).

Gastruloids were fixed and stained as previously described^10^. Briefly, gastruloids were fixed overnight in 4% paraformaldehyde at 4°C. The following day, they were washed 3 times for 1 hour each with PBST and incubated in blocking buffer (PBS containing 0.1% BSA and 0.3% Triton X-100) overnight at 4°C. Primary antibodies were then applied, diluted in blocking buffer to working concentrations as per manufacturer’s recommendations, and incubated overnight at 4 °C. Stained gastruloids were washed with washing buffer (PBS containing 0.3% Triton X-100), stained with secondary antibodies (diluted either 1:500 or 1:1000 in blocking buffer) and DAPI (diluted 1:1000) overnight at 4 °C in the dark. The following day gastruloids were washed in blocking buffer and mounted in SlowFade™ Gold Antifade Mountant (S36936, Thermo Fisher Scientific). The antibodies used in this study are listed in Supplementary Table 14. All samples were analyzed with the Nikon Eclipse Ti2 confocal microscope and analyzed using FIJI^141^ and the python sci-kit-image^142^.

### RNA-seq analysis

#### Sample preparation

Each stage consisted of 2 biological replicates harvested within the same experimental batch to minimize batch effects. Approximately 0.5 million cells per replicate were harvested across mouse and human cells across the 4 gastruloid developmental stages. DNA and RNA from each sample were isolated using the Qiagen AllPrep DNA/RNA kit (Qiagen #80204). Approximately, 500ng of total RNA was used as input for library preparation. mRNAs were isolated using the NEBNext Poly(a) mRNA Magnetic Isolation Module (NEB #E7490) and prepared for sequencing using the NEBNext UltraII RNA Library Prep Kit for Illumina (NEB #E7770).

#### Sequencing and data analysis

Concentrations of cDNA libraries across all samples were estimated from either the Qubit (Invitrogen) and/or visualized by Tapestation (Agilent) to ensure standard ranges for library sizes. All libraries were dual-indexed with 8 nucleotide indexes using NEBNext® Multiplex Oligos for Illumina® (Index Primers Set 1) and were sequenced on NextSeq 2000 (Illumina) either by 2×150bp or 2×50bp configuration.

Basecall files were converted to fastq formats using bcl2fastq (Illumina) and demultiplexed on the i5 and i7 indexes. FastQC was performed to estimate the quality of the reads. Adapter trimming and filtering for low-quality reads was performed using Trimmomatic v0.39^143^, either in paired-end or single-end mode, trimming low-quality reads (<2) at the ends and applying a four base sliding window across reads, retaining those with average quality above 15. Depending on the species, trimmed reads were then aligned using STAR^144^ to either the human GRCh38 or mouse GRCm39 reference assemblies. Human samples had an average unique mapping rate of 64% while those of mouse samples were 48%. Finally, count matrices for each species were then generated with the bam files using FeatureCounts.

### Mass spectrometry data collection

#### Sample preparation

For each stage analyzed, we collected 1-2.5 million cells per replicate across four gastruloid developmental stages. To mitigate batch effects, all replicates from each developmental time point were harvested together within the same batch. Stem cells across each stage were harvested from culture plates by enzymatic dissociation using Accutase^TM^ (StemCell Technologies, #07920). Since each gastruloid is cultured in a single well of a 96-well U-bottom plate, gastruloids were first pooled together to reach the 2.5 million cell number and gently centrifuged at 500g for 5 min to remove growth media followed by Accutase treatment to dissociate the gastruloids. Once dissociated, Accutase treatment for both gastruloid and stem cell samples was quenched by addition of a wash buffer consisting of either StemFlex or mTeSR+ along with rock inhibitor (Y-27632). Finally cells were washed twice with PBS to remove cell debris, lysed cells, and matrigel from the samples. Samples were finally stored at -80C after aspirating the PBS before proceeding to protein isolation.

Cell pellets were thawed on ice and resuspended in lysis buffer (8M urea, 250mM EPPS pH 8.5, 50 mM NaCl, Roche protease inhibitor cocktail, Roche PhosSTOP). The cell pellets were homogenized using a 21-gauge needle to syringe pump lysate. Lysates were cleared by centrifugation at 21,130 g at 4°C for 30 minutes. Supernatants were placed in clean microcentrifuge tubes and a BCA assay (Pierce) was performed to determine protein concentrations. Lysate containing 25 ug of protein material for biological triplicates at each point of gastrulation were reduced and alkylated with 5 mM Dithiothreitol (DTT) for 30 minutes at room temperature and 20 mM Iodoacetamide (IAA) for 1 hour in the dark at room temperature. The IAA reaction was then quenched with 15 mM DTT. Single-pot solid phase sample preparation (SP3)^145^ using Sera-Mag SpeedBeads was performed to desalt the reduced and alkylated samples. An on-bead protein digestion was performed by adding LysC at a 1:100 ratio (protease:protein) overnight (16-24 hours) on a thermocycler at room temperature then adding trypsin at a 1:100 ratio for 6 hours at 37°C at 900 rpm. TMTpro was used to label each sample at a 2.5:1 ratio of TMTpro reagents to the peptide mixtures for each sample. Samples were left at room temperature for 1 hour for TMTpro labeling and labeling efficiency was verified to be >99% for lysines and >97% for N-termini. The labeling reaction was quenched with 5% hydroxylamine diluted to a concentration of 0.3% for 15 minutes at room temperature. Samples were then placed on a magnetic rack to aggregate SP3 beads and labeled peptide supernatants from each sample were pooled. The pooled sample was then partially dried down by speed-vac and 10% formic acid was added to bring the pH of the pooled sample to below 3 for desalting. The pooled sample was desalted using a Sep-Pak C18 cartridge (Waters) and then dried down completely.

#### Phosphoproteomics sample preparation

Pooled sample was resuspended in 94 uL of 80% acetonitrile and 0.1% trifluoroacetic acid for Fe^3+^-NTA magnetic bead phosphopeptide enrichment^146^. 100 uL of 75% acetonitrile 10% formic acid was added to a clean microcentrifuge tube and the Fe^3+^-NTA magnetic beads were washed twice with 1 mL of 80% acetonitrile and 0.1% trifluoroacetic acid and the supernatant was removed. After the final wash, the peptides in 94 uL of 80% acetonitrile and 0.1% trifluoroacetic acid were added to the tube with the washed beads. The sample was vortexed and incubated for 30 min on thermoshaker (250 rpm, 25 C). After the incubation period, the sample was washed 3 times with 200 uL of 80% acetonitrile and 0.1% trifluoroacetic acid and all flowthrough was saved in a clean microcentrifuge tube as it contains non-phosphorylated peptides. 100 uL of 50% acetonitrile and 2.5% NH_4_OH was added to elute phosphorylated peptides from magnetic beads and then sample was transferred to tube with 100 uL of 75% acetonitrile and 10% formic acid. The phosphopeptide enriched sample was dried down immediately by speed-vac and resuspended in 100 uL of 5% formic acid and a C18 stage tip was used to desalt the phosphopeptide enriched sample. The sample was then transferred to a MS insert vial that was placed within a microcentrifuge tube. The sample was placed in a -80 C freezer for 30 minutes and then dried down completely in a speed vacuum. The sample was then resuspended in 10 uL of 2% formic acid and 5% acetonitrile within the MS insert vial.

#### Total proteomics sample preparation

The saved flowthrough was dried down using the speed vacuum, resuspended in 500 uL of 5% formic acid, and a Sep-Pak C18 cartridge (Waters) was used to desalt the sample. The flowthrough sample was dried down completely in speed vacuum after desalting. The flowthrough sample was resuspended and neutralized in 1 mL of 10 mM ammonium bicarbonate/90% acetonitrile and dried down completely in speed vacuum again. Sample was resuspended in 115 uL 10 mM ammonium bicarbonate and 5% acetonitrile and 110 uL were transferred to a sample vial. High-pH Reverse-Phase HPLC Fractionation was performed on the flowthrough sample using an Agilent 1200 HPLC system. After HPLC fractionation^147^, fractions were dried down in speed-vac, resuspended in 100 uL of 5% formic acid, and cleaned via C18 stage tip. Elution from each stage tipped fraction was placed in a MS insert vial and dried down in vial. Fractions were then resuspended in 5 uL of 2% formic acid 5% acetonitrile within the MS insert vial.

### Mass spectrometry data acquisition

#### Proteomics

All analyses were performed using an Orbitrap Eclipse Tribrid Mass Spectrometer (Thermo Fisher Scientific) in-line with an Easy-nLC 1200 autosampler (Thermo Fisher Scientific). The peptides underwent separation using a 15 cm-long C18 column with a 75 μm inner diameter, with a particle size of 1.7 μm (IonOpticks). Each fraction collected from the off-line fractionation was analyzed using a 90 min gradient of 2% to 26% acetonitrile in 0.125% formic acid with a flow rate of 500 nl/min. The MS1 resolution was set to 120,000 with a scan range of 400-2000 m/z, a normalized automatic gain control (AGC) target of 200%, and a maximum injection time of 50 ms. The FAIMS voltage was cycled between activated at a constant compensation voltages (CV) of -40 V, -60, and -80 V. MS2 scans were collected with an AGC target of 200%, maximum injection time of 50 ms, isolation window of 0.5 m/z, CID collision energy of 35% (10ms activation time), and “Rapid” scan rate. SPS-MS3^148^ scans were triggered based on the real-time search (RTS) filter^36^. Briefly, RTS was run by searching species specific Uniprot protein databases (downloaded 04/2023) for mouse (taxid: 10090) and human (taxid: 9606) with static modifications for carbamidomethylation (57.0215) on cysteines and TMTpro acylation (304.2071) on peptide N-termini and lysines; variable modification of oxidation (15.9949) on methionines, one missed cleavage, a maximum of three variable modifications per peptide. Scan parameters of the SPS-MS3 were set to collect data on 10 SPS ions at a resolution of 50,000, AGC target of 400%, maximum injection time of 150 ms, and HCD normalized collision energy of 45%.

#### Phosphoproteomics

Duplicate injections (4 μL) were analyzed on an Orbitrap Eclipse Tribrid Mass Spectrometer (Thermo Fisher Scientific) along with an Easy-nLC 1200 autosampler (Thermo Fisher Scientific). The peptides underwent separation using a 15 cm-long C18 column with a 75 μm inner diameter, with a particle size of 1.7 μm (IonOpticks). Each fraction was analyzed using a 90 min gradient of 2% to 26% acetonitrile in 0.125% formic acid with a flow rate of 400 nl/min. The MS1 scan resolution was set to 120,000 with a scan range of 400-1800 m/z, a normalized AGC target of 200%, and a maximum injection time of 50 ms. The FAIMS voltage was cycled between compensation voltages of -40, -60, and -80 V. MS2 scans were collected with an AGC target of 250%, maximum injection time of 35 ms, isolation window of 0.5 m/z, CID-Multistage Activation (MSA) collision energy of 35% (10ms activation time) with additional activation of the neutral loss mass of n-97.9763, and “Rapid” scan rate. For SPS-MS3 scans^148^ a resolution of 50,000, AGC target of 300%, maximum injection time of 86 ms, and HCD normalized collision energy of 45%.

### Proteomic and phosphoproteomic data analysis

#### Peptide spectral matching

Raw files were searched against the relevant annotated proteome from Uniprot (Human: October 2020; Mouse: March 2021). Sequences of common contaminant proteins and decoy proteins were added to the Uniprot FASTA file to also be searched. Comet search algorithm^149^ was utilized to match peptides to spectra with the following parameters: 20 ppm precursor tolerance, fragment_tolerance of 1.005, TMTpro labels (304.207145) on peptide N-termini and lysine residues, alkylation of cysteine residues (57.0214637236) as static modifications, and methionine oxidation (15.9949146221) as a variable modification. Phosphoproteomics runs were also searched for phosphorylation as a variable modification on serine, threonine, and tyrosine residues (79.9663304104). Peptide-spectrum matches (PSMs) were filtered to a 1% false discovery rate (FDR) using a linear discriminant analysis^36^. Proteins were filtered to an FDR of 1% using the rules of protein parsimony and the protein picker methods^150^. For quantitation, PSMs were required to have a summed TMT reporter ion signal-to-noise ≥100^148^.

#### Differential protein expression analysis

Differentially expressed proteins (DEPs) between developmental gastruloid stages were identified as follows. For each protein, we calculated the log2 ratios of mean abundance across 2 given timepoints and computed their p-values using a standard t-test. We corrected for multiple hypothesis testing by adjusting the p-values using the Benjamini-Hochberg (BH) procedure. We classified proteins as DEPs if they had an absolute fold change of greater than between the 2 and BH-adjusted p-value < 0.05 between 2 given timepoints.

#### Protein module analysis

All quantified proteins were mapped onto known transcription factors (curated from the Transcription Factor Database^151^, protein complexes (curated from CORUM^64^ and EMBL ComplexPortal^65^), biochemical pathways (curated from BioCarta^59^, KEGG^60^, PID^62^, Reactome^61^ and WikiPathways^58^), subcellular localization (curated from Human Protein Atlas^30,152^), and Gene Ontology (GO) terms^42^. For biochemical pathways and complexes, we filtered module sets to those where we detected greater than 2 members. With respect to subcellular locations, if a protein in Human Protein Atlas was listed as localized to multiple regions in its main subcellular location, we considered each location as unique. We avoided searching our data against overly broad descriptions of GO terms by filtering for terms containing fewer than or equal to 150 genes and greater than 2 members detected in our data. All mappings were based on Uniprot annotations^71,153^ unless otherwise stated.

#### Correlation network construction and network analysis

We first intersected the human and mouse protein datasets and used 6,261 proteins that were observed across the shared timepoints within a cell line *i.e.* primed ESCs, early and late gastruloids. We normalized each protein’s abundance in given replicate to its respective species geometric mean and log_2_ transformed values for subsequent analysis unless otherwise stated. To construct our correlation network, we first calculated the Pearson correlation coefficients (r_Pearson_) across all 19,596,930 possible pairs of proteins. Since we already calculated r_Pearson_ across all possible pairs of proteins, we permuted sample labels across our dataset to generate the null distribution of correlation coefficients. We then stringently filtered the network edges with Benjamini-Hochberg (BH) adjusted p-values < 0.01 and absolute r_Pearson_ >= 0.95. This step filtered the network down to 489,417 (301,561 correlated and 187,856 anticorrelated) pairs and was used for subsequent network analysis.

#### Edge annotation in correlation network

We considered seven major annotations as literature evidence for any given edge: 1) Protein-protein interaction, 2) Belonging to the same protein complex or 3) biochemical pathway, 4) GO biological process, 5) GO molecular function, 6) GO cellular component or 7) Subcellular location. Protein complex annotations were obtained from CORUM^64^ (downloaded 9/12/2022) and ComplexPortal^65^ (downloaded 1/7/2024). Annotated gene sets for pathways^58–62^ and GO^42^ were downloaded from the Molecular Signatures Database^154^. Protein localization annotations were curated from Human Protein Atlas^30,152^. Networks were illustrated using the igraph R package or Cytoscape^155^.

#### Bioinformatic identification cooperative protein interactions

We searched all nodes in our correlation network against known complexes and pathways which consisted of at least 3 subunits. We adapted a previously described approach^63^ and employed a Fisher’s exact test to compute statistical enrichment of cooperative complexes with established modules. For each protein complex or pathway module, we tested its neighboring proteins (first-degree edges) for significant association with a particular module and termed those as cooperative proteins. For each protein tested, we first counted the number of edges that it shared with the established module, second we counted the number of edges that linked the module to other proteins (excluding the candidate protein) in the network, third we counted the number of edges the candidate protein had to rest of the correlation network (*i.e.* excluding the module of interest) and finally, we counted the number of edges that were not associated with the candidate protein nor the module of interest. These edge counts were used to compute statistical significance using Fisher’s exact test. We independently repeated this test for all 6,261 proteins against 1,357 known protein complexes and select metabolic pathways. The p-values obtained were adjusted for multiple hypothesis testing using the BH procedure and only cooperative proteins with adjusted p-value < 0.05 were considered significant.

#### Comparison of RNA and protein abundance analysis

Global RNA-protein correlations were calculated using all 9 observations of transcripts and proteins across mouse and human gastruloid development. To ensure stringent analysis, we filtered for genes detected in both species for the downstream analysis. Pseudocounts of 1 were added to filtered count matrices and were converted to transcripts per million (TPM). Mean transcript and protein abundances were converted to log2 fold change ratios to their respective species geometric mean. For every gene, we calculated the per-gene RNA-protein correlation (r_Pearson_) using a vector of abundances across 9 samples. GO term enrichment of biological processes in correlelated and anticorrelated genes was performed using ClusterProfiler^156^. We intersected the 6010 genes detected across both datasets with Human Protein Atlas^30^ for subcellular locations, CORUM^64^ and ComplexPortal^65^ for protein complexes and KEGG for biochemical pathways^60^. To measure the extent of correlation of transcripts and RNAs within mouse timepoints we calculated the ratio of protein to RNA mean fold changes across each timepoint. In summary, a discordance of 0 implied that the protein and RNAs were highly correlated while discordance less than 0 implied that the RNAs were more abundant than protein levels and vice versa. Discordance scores for protein complexes was calculated by taking the median protein-RNA correlation across constituent members. To prevent averaging pairs of proteins, we only considered complexes where more than 2 proteins were detected in our data. Transcriptional signatures of stage specific mouse transcription factors were detected as follows. First, we calculated the Pearson correlation comparing transcription factor protein abundances to all observed transcripts. We subset the resulting correlation matrix to identify protein-transcript pairs with high correlation (r_Pearson_ >= 0.9) and used TFLink^83^ to select only transcripts that were annotated as targets of specific transcription factors. We confirmed the identified transcription factor targets displayed similar temporal regulation to their upstream transcription factor by comparing target transcript abundance at each stage to determine the maximum transcript abundance.

#### Phosphoprotein and Kinase analysis

For differential expression testing and analysis, in every pairwise comparison, log2 ratios for all quantified phosphosites were calculated following subtraction of the log2 ratios of the corresponding proteins to identify protein independent phosphorylation changes. Kinase substrate pairs were curated from PhosphositePlus^90^. Human kinases were annotated using KinMap^103^. For Kinase-Substrate prediction and enrichment analysis, for each phosphosite, we first calculated the log2 fold change ratio to the row mean (across all samples) subtracted the corresponding protein log2 fold change ratios and used that as input into the KSEA app^89^ with a minimum substrate cutoff >=2 to calculate z-scores for kinases. Kinase substrate pairs with absolute r_Pearson_ >= 0.5 were visualized as a network using Cytoscape^155^.

## ACKNOWLEDGMENTS

At the University of Washington, the authors thank Diego Calderon, Chengxiang Qiu, Jean-Benoît Lalanne, Aidan Keith, and, Shawn Fayer along with the rest of the members in the Shendure and Starita labs particularly for critical insights, discussions, and feedback. The authors thank Valentino Browning, Eva Nichols, and Katie Partington for assistance and advice related to microscopy and imaging. The authors also thank Akshaya Rajaraman and Kevin Drew (University of Illinois Chicago) for advice and feedback related to network analyses and mapping protein complexes.

R.K.G. acknowledges support from the Washington Research Foundation postdoctoral fellowship. D.K.S. acknowledges support from the NIH/NIGMS (R35GM150919), Washington Research Foundation, the W.M. Keck Foundation, an Andy Hill CARE Distinguished Researcher Award, a Cancer Consortium New Investigator Award, and the Pew Charitable Trusts. R.K.G., S.C., S.B., L.M.S. were supported by the National Human Genome Research Institute (NHGRI; 1RM1HG010461). J.S. is an Investigator of the Howard Hughes Medical Institute and acknowledges support from the Paul G. Allen Frontiers Group (Allen Discovery Center for Cell Lineage Tracing) and the Brotman Baty Institute for Precision Medicine.

## AUTHOR CONTRIBUTIONS

R.K.G. and N.H. in consultation with D.K.S. conceived the study. R.K.G. and N.H. performed stem cell and gastruloid experiments. R.K.G. and N.H. performed the transcriptomics experiments with assistance from S.C. and S.B.. V.L., R.F., and C.D.M. performed the proteomics and phosphoproteomics experiments. R.K.G. computationally analyzed the data with support from N.H., M.S., D.K.S.. J.S. with assistance from R.K.G built the web-interface. R.K.G., N.H., V.L., D.K.S., L.M.S., and J.S., wrote the manuscript. D.K.S., N.H., L.M.S., and J.S oversaw the experiments and data analyses.

## COMPETING INTERESTS

J.S. is a scientific advisory board member, consultant and/or co-founder of Cajal Neuroscience, Guardant Health, Maze Therapeutics, Camp4 Therapeutics, Phase Genomics, Adaptive Biotechnologies, Scale Biosciences, Prime Medicine, Somite Therapeutics, Sixth Street Capital and Pacific Biosciences. D.K.S. is a consultant and/or collaborator with ThermoFisher Scientific, AI Proteins, Genentech, and Matchpoint Therapeutics. The other authors declare no competing interests.

## SUPPLEMENTARY TABLES

Supplementary Table 1- Quantified protein intensities across human and mouse gastruloid development datasets

Supplementary Table 2- Gene Ontology (GO) enrichments of protein clusters with similar temporal profiles

Supplementary Table 3- Pairwise protein correlation network

Supplementary Table 4- Summary statistics of the protein correlation network

Supplementary Table 5- Cooperative proteins of protein complexes

Supplementary Table 6- Jaccard index matrix of cooperative protein pair overlap

Supplementary Table 7- Protein-RNA discordance across the entire dataset

Supplementary Table 8- Protein-RNA correlation across protein complexes and pathways

Supplementary Table 9- Stage-specific GO enrichments for discordant gene sets across mouse gastruloid development

Supplementary Table 10- Quantified phosphosite intensities across human and mouse gastruloid development datasets

Supplementary Table 11- Phosphosites of proteins downstream of pluripotency markers POU5F1, NANOG, and POU5F1

Supplementary Table 12- Disease associated genes and complexes quantified in the human dataset

Supplementary Table 13- gRNA sequences used for RNP-based perturbation of Commander co-regulatory network

Supplementary Table 14- Antibodies used in this study

## DATA AND CODE AVAILABILITY

RNA-seq data have been deposited to the Gene Expression Omnibus (GEO) database with the identifier GSE273813. Mass spectrometry proteomics data have been deposited to the ProteomeXchange Consortium^157^ via the PRIDE partner repository^158^ with the dataset identifier PXD054460. Reviewers can access these data through PRIDE using the account details: Username; Password: 4lQJ5v6pqvGs.

All supporting scripts and code have been deposited onto the following repository at https://github.com/bbi-lab/Temporal-Gastrulomics. All processed data are available through the web application at: https://gastruloid.brotmanbaty.org/.

## Notes

### Summary of Updates

Figures and text in the manuscript updated to reflect new results. Supplemental files updated

